# Insertases Scramble Lipids: Molecular Simulations of MTCH2

**DOI:** 10.1101/2023.08.14.553169

**Authors:** Ladislav Bartoš, Anant K. Menon, Robert Vácha

## Abstract

Scramblases play a pivotal role in facilitating bidirectional lipid transport across cell membranes, thereby influencing lipid metabolism, membrane homeostasis, and cellular signaling. MTCH2, a mitochondrial outer membrane protein insertase, has a membrane-spanning hydrophilic groove resembling those that form the lipid transit pathway in known scramblases. Employing both coarse-grained and atomistic molecular dynamics simulations, we show that MTCH2 significantly reduces the free energy barrier for lipid movement along the groove and therefore can indeed function as a scramblase. Notably, the scrambling rate of MTCH2 *in silico* is similar to that of VDAC, a recently discovered scramblase of the outer mitochondrial membrane, suggesting a potential complementary physiological role for these mitochondrial proteins. Finally, our findings suggest that other insertases which possess a hydrophilic path across the membrane like MTCH2, can also function as scramblases.

**Graphical abstract:** 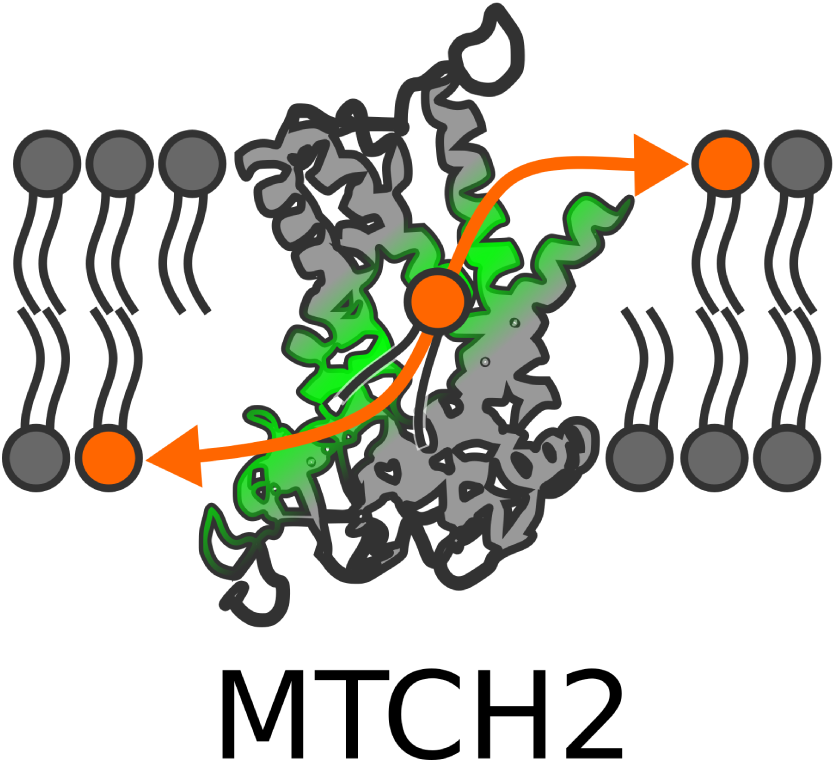

**Highlights:**

- Scrambling activity of MTCH2 identified using computer simulations
- MTCH2 may act redundantly with VDAC as outer mitochondrial membrane scram-blase
- Insertases and scramblases may share a common functional mechanism

## Introduction

The spontaneous movement of polar lipids from one side of a membrane bilayer to the other is slow and is accelerated in cells to a physiologically relevant rate by lipid transporters^1,2^. Flippases and floppases are transporters which move lipids unidirectionally, against their concentration gradient in an ATP-dependent manner, whereas scramblases enable bidirectional lipid movement without need of metabolic energy. Scramblases typically possess transmembrane helices that are not fully hydrophobic. These helices form a membrane-spanning hydrophilic groove along which phospholipids can move in analogy to a credit card (phospholipid) being swiped through a card reader (scramblase) ^1^. Known scramblases are alpha-helical proteins^3–6^, but recently, a beta-barrel scramblase, the voltage-dependent anion channel (VDAC), was identified^7^. The VDAC barrel has membrane-facing polar residues that, when paired with the same residues in a second beta barrel at a dimer interface, create a local membrane defect along which lipids cross the membrane in credit card mode. Therefore, the presence or ability to form a hydrophilic groove across the membrane seems to be a common feature for lipid scrambling.

Insertases assist in the integration of membrane proteins into lipid bilayers^8^. Their activity is necessary because membrane proteins typically have hydrophilic segments that do not spontaneously insert into the membrane. The translocating moieties of these proteins are thus amphiphilic, similar to lipids, and it is interesting to consider whether insertases and scramblases may share a common mechanism even though lipids and their polar headgroups are generally smaller than the translocating parts of membrane proteins. In fact, it has been suggested that scramblases, insertases, and some translocases may facilitate the insertion and translocation of phospholipids and proteins through membrane thinning^9^.

Recent work demonstrated that the mitochondrial outer membrane protein MTCH2 is an insertase^10^. MTCH2 has a wide transbilayer groove lined with polar and charged residues, reminiscent of the hydrophilic grooves of scramblases, leading us to hypothesize that it may function as a scramblase. To test this hypothesis, we performed coarse-grained and atomistic molecular dynamics (MD) simulations of MTCH2 in a model lipid bilayer. We show that MTCH2 is indeed a scramblase, with the hydrophilic groove providing a transbilayer path for the movement of lipid headgroups while the acyl chains move through the hydrophobic interior of the membrane in credit card mode.

## Results and discussion

We used a MTCH2 structure predicted by AlphaFold^11,12^ and simulated it in 1-palmitoyl-2-oleoyl-sn-glycero-3-phosphocholine (POPC) membrane to observe its potential scrambling activity.

We started with coarse-grained simulations using the Martini 3 force field^13^. We conducted three unbiased simulations (each of 10 *μ*s), to analyze the spontaneous scrambling activity of MTCH2. Additionally, we calculated the free energy of lipid flip-flop along the MTCH2 groove and compared it with the free energy of flip-flop in a proteinless POPC membrane.

In the unbiased simulations with MTCH2, we observed rapid insertion of lipids from the upper leaflet into the cavity of MTCH2 (see Figure 1 A-B). The cavity of MTCH2 is situated between α-helices 1 (residues 1–31), 3 (residues 74–100), 4 (residues 117–151), 6 (residues 182–207), and 7 (residues 216–247). It is shaped like a funnel that opens towards the leaflet with the N-terminus and is lined with a large number of hydrophilic and charged amino acid sidechains (see Figure 1 B). This cavity has already been described as the structural basis of the insertase activity of MTCH2 by Guna et al^10^.

**Figure 1:**
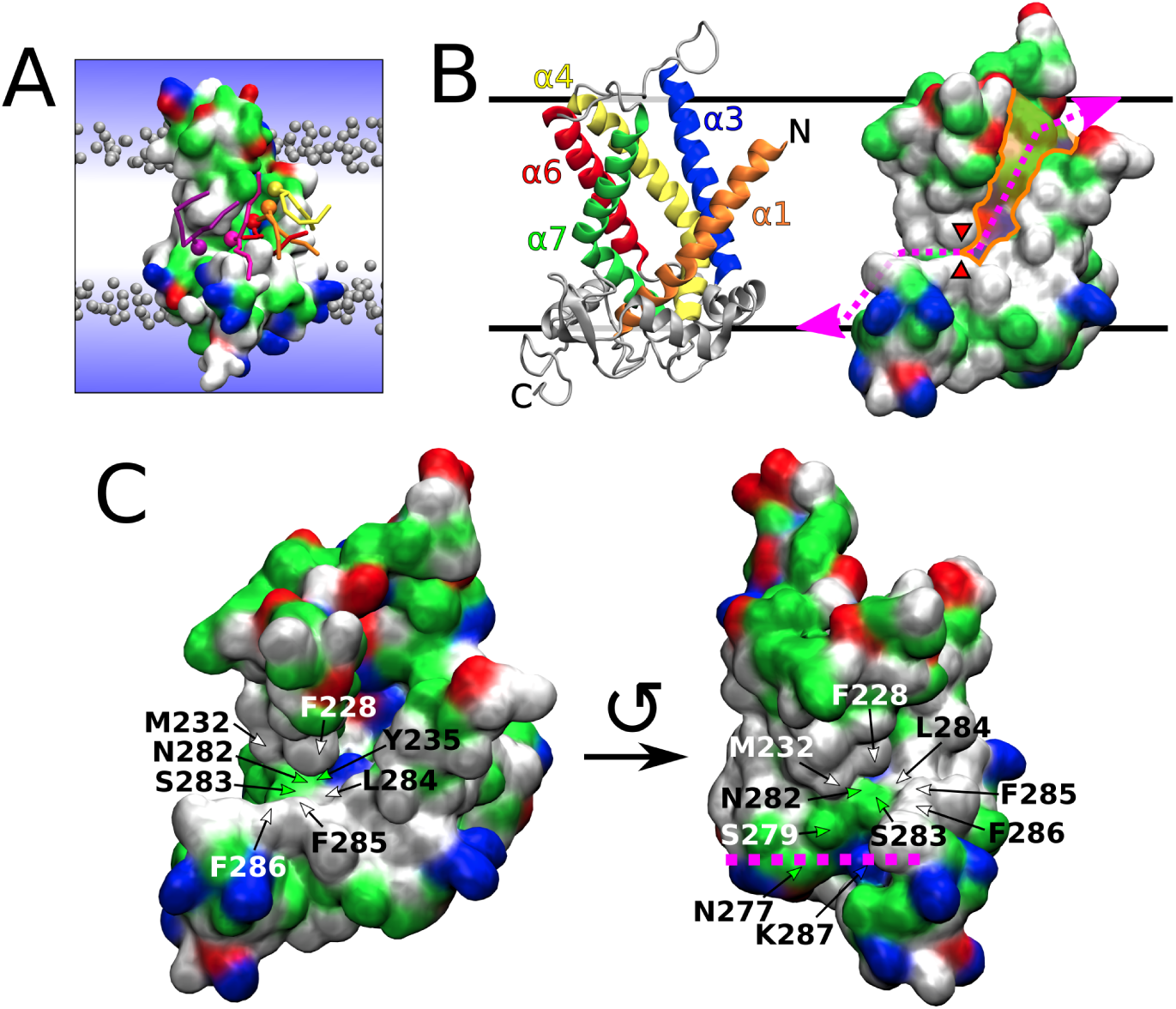
Scrambling pathway of MTCH2. A) Snapshot from a Martini 3 simulation showing several lipids (colored in yellow, orange, red, magenta, and purple) translocating along the scrambling pathway of MTCH2. The protein is shown as a molecular surface colored according to the character of its residues (hydrophilic = green, hydrophobic = white, positively charged = blue, negatively charged = red). Bulk lipids are depicted as gray beads (phosphate groups) with hydrophobic tails omitted for clarity. Water is represented only schematically as a blue gradient, omitting the presence of water in the cavity of the protein for visual clarity. B) Two visualisations of the atomistic structure of MTCH2 after 3 *μ*s of atomistic simulation. Left: Cartoon representation showing α-helices forming the cavity of the protein. Right: Molecular surface representation showing the cavity (colored in orange), the scrambling pathway (dotted magenta line) and the position of the “hydrophobic gate” (red wedges). C) More detailed views of the atomistic structure of MTCH2 showing residues of the C-terminal pathway and the “hydrophobic gate”. The structure on the left depicts the frontal and slightly superior view of the MTCH2 cavity. The structure on the right shows MTCH2 rotated counterclockwise by roughly 90*^◦^* showing frontal view of the C-terminal pathway, while the cavity is mostly hidden. Purple line shows the approximate position of the lipid headgroups in the lower leaflet. The top of the figure represents the cytosol, and the bottom represents the intermembrane space of mitochondria.

In our Martini 3 simulations, the insertion of lipids into the cavity was quickly followed by a large number of flip-flop events during which lipids traversed between the membrane leaflets in both directions (see Figure 2 A Left). We identified a mostly hydrophilic pathway extending the cavity towards the opposite leaflet (with C-terminus) and allowing lipids to pass along MTCH2 through the entire hydrophobic membrane core. This extended hydrophilic pathway (further called “C-terminal pathway”) starts with Y235 located in helix 7 near the bottom of the cavity and continues with N282, S283, S279, K287, and N277 until the phosphate level of the lower membrane leaflet is reached. The entry into this pathway is however partly obstructed by several hydrophobic residues (further called “hydrophobic gate”), namely F228, M232 (*α*-helix 7) and L284, F285, and F286 (all located in a C-terminal loop following *α*-helix 8). See Figure 1 C for a more detailed view of the C-terminal pathway and the “hydrophobic gate”. Also see Figure S1 for simulation snapshots showing membrane and lipid structure near the cavity of MTCH2.

**Figure 2:**
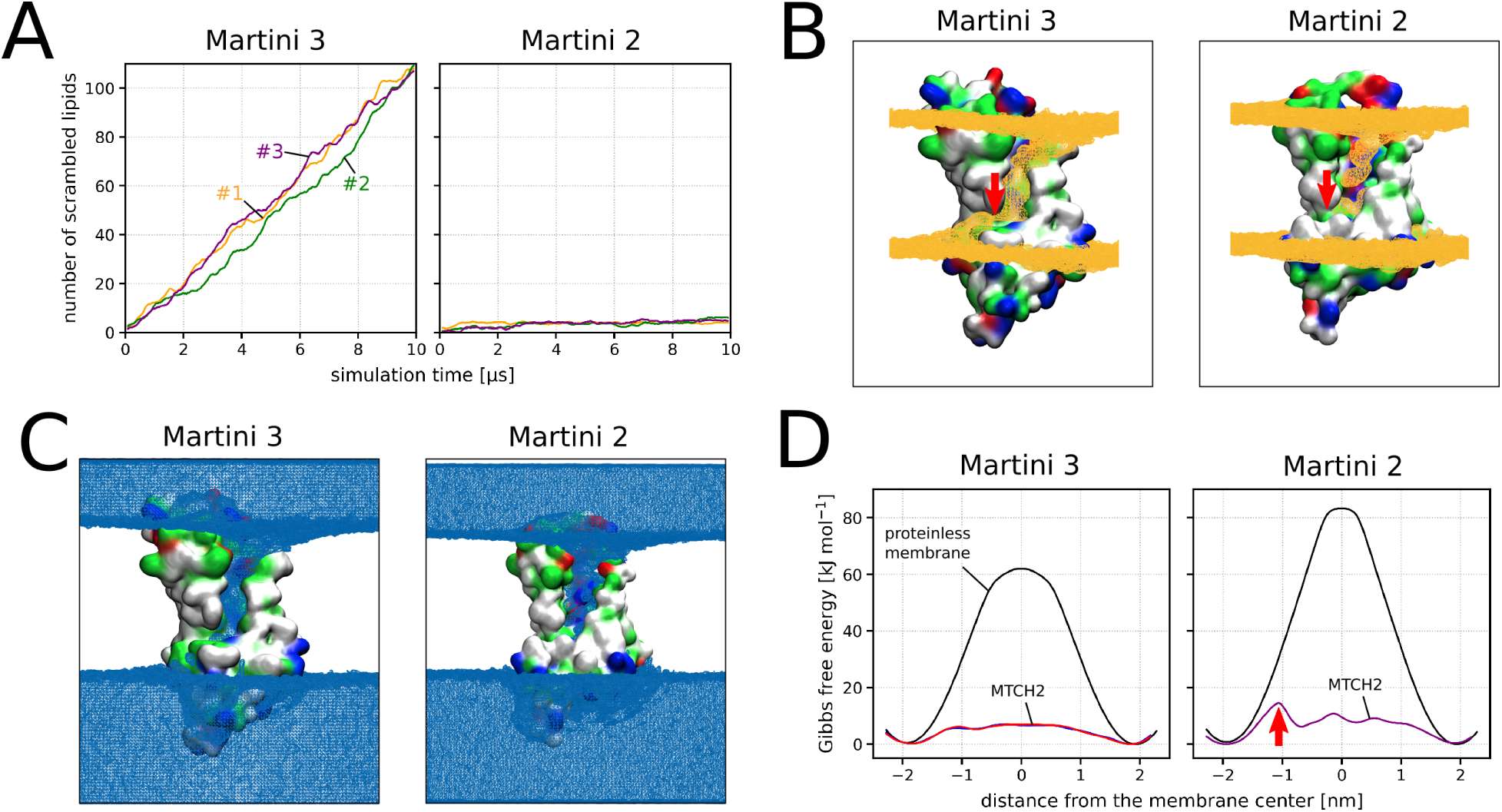
Comparison of Martini 3 and Martini 2 simulations. A) Scrambling rate of MTCH2 in a POPC membrane shown as the number of “scrambled lipids” in time (running average over 200 ns). Each line corresponds to single 10 *μ*s long Martini simulation. Lipid was considered to be scrambled when it was located in the opposite leaflet to its original one. Scrambling rate of lipids in a proteinless POPC membrane is zero in both models (on the simulated time scale). B) Density of phosphate beads around MTCH2 depicted as orange isosurface wireframe. Lipid phosphates readily insert into the cavity of MTCH2 in both models. In Martini 3, they are also able to easily enter the C-terminal pathway, while this is not the case for Martini 2. The position of the “hydrophobic gate” is indicated by the red arrow. C) Density of water beads around MTCH2 depicted as blue isosurface wireframe. In both models, a large number of molecules of water inserts into the cavity of MTCH2. D) Free energy profiles of lipid flip-flop in a proteinless POPC membrane (black) and along the MTCH2 scrambling pathway (red, blue, and purple). In both models, the presence of MTCH2 dramatically reduces the flip-flop free energy barrier. The red arrow in the Martini 2 chart indicates a peak in the free energy profile corresponding to the “hydrophobic gate”. The calculation error is < 1 kJ mol*^−^*^1^. See Figure S3 for detailed view of free energy calculations convergence.

To further support our qualitative observations and demonstrate local membrane defect/thinning, we calculated the density of phosphates and the density of water around MTCH2 in all three unbiased simulations. As shown in Figure 2B Left, the density of phosphates is significantly increased in the cavity of the protein as well as in the C-terminal pathway indicating that phosphates readily pass through the membrane along MTCH2. The membrane is also dramatically thinned in the cavity, as shown in Figure S2. Similarly, Figure 2C Left shows the increased density of water in the cavity of MTCH2. Unlike phosphates, water density is reduced (but not absent) along the C-terminal pathway.

The identified features of MTCH2, including the hydrophilic pathway, water defect, and membrane disruption, align with those observed in other scramblases, such as TMEM16^3,14^, opsin^4,15^, and VDAC^7^. MTCH2 shares with the other scramblases the “credit card mechanism” of lipid scrambling, which involves the lipid head contacting a hydrophilic groove of the protein while the lipid tails face the hydrophobic core of the membrane^1^.

To provide a more quantitative picture of the scrambling activity of MTCH2, we calculated the free energy of the entire flip-flop process along the identified scrambling pathway. As shown in Figure 2D Left, the presence of MTCH2 dramatically decreased the free energy barrier for lipid flip-flop by almost 90% compared to flip-flop in proteinless POPC bilayer (from 62 to just 7 kJ mol*^−^*^1^). See Figures S4 and S5 for representative snapshots from umbrella sampling windows used to calculate the free energy profiles.

As Martini 3 may underestimate the free energy barrier for the flip-flop process and consequently overestimate the flip-flop rate, we also performed the same simulations using the older Martini 2.2 force field^16–18^. Despite observing a high tendency of lipids and water to enter the cavity of MTCH2, we only observed a handful of complete flip-flop events within the simulated timescale (Figure 2A Right) likely due to the hindrance of the “hydrophobic gate” described above. Based on these results, we speculate that the opening and closing of the “hydrophobic gate” could regulate the scrambling activity of MTCH2.

As with Martini 3, we calculated the density of phosphates (Figure 2B Right) and water (Figure 2C Right) around MTCH2 showing the tendency of both phosphates and water to enter the cavity of the protein as well as the inability of phosphates to finish the C-terminal part of the scrambling pathway.Nevertheless, as seen from the calculated free energy profiles (Figure 2D Right), the presence of MTCH2 still decreases the free energy barrier of the flip-flop process by about 80% compared to the proteinless membrane (from 83 to 15 kJ mol*^−^*^1^). See Figure S6 for additional information about the Martini 2 free energy calculations and selected snapshots from the process.

We acknowledge that the MTCH2 structure predicted by AlphaFold may have some limitations. For instance, the stability of the structure in the membrane may change and the predicted cavity could collapse or only open under specific conditions. In our coarse-grained simulations, the secondary and tertiary structure of MTCH2 were restrained, thereby limiting their dynamics and preventing us from evaluating the stability of the predicted structure. To address this limitation, we simulated MTCH2 using the atomistic CHARMM36m force field^19^ in a POPC membrane. In a 3 *μ*s simulation at physiological conditions, we observed no significant changes in the structure apart from repositioning of the loops (Figure 3A,B), suggesting the stability of the predicted structure. To further validate this stability, we performed an additional 1.5 *μ*s simulation of MTCH2 at 330 K (57 *^◦^*C), which increases susceptibility to structural alterations and no structural changes were observed. Our observations indicate that the predicted structure is stable and the cavity remains accessible to lipids and water.

**Figure 3:**
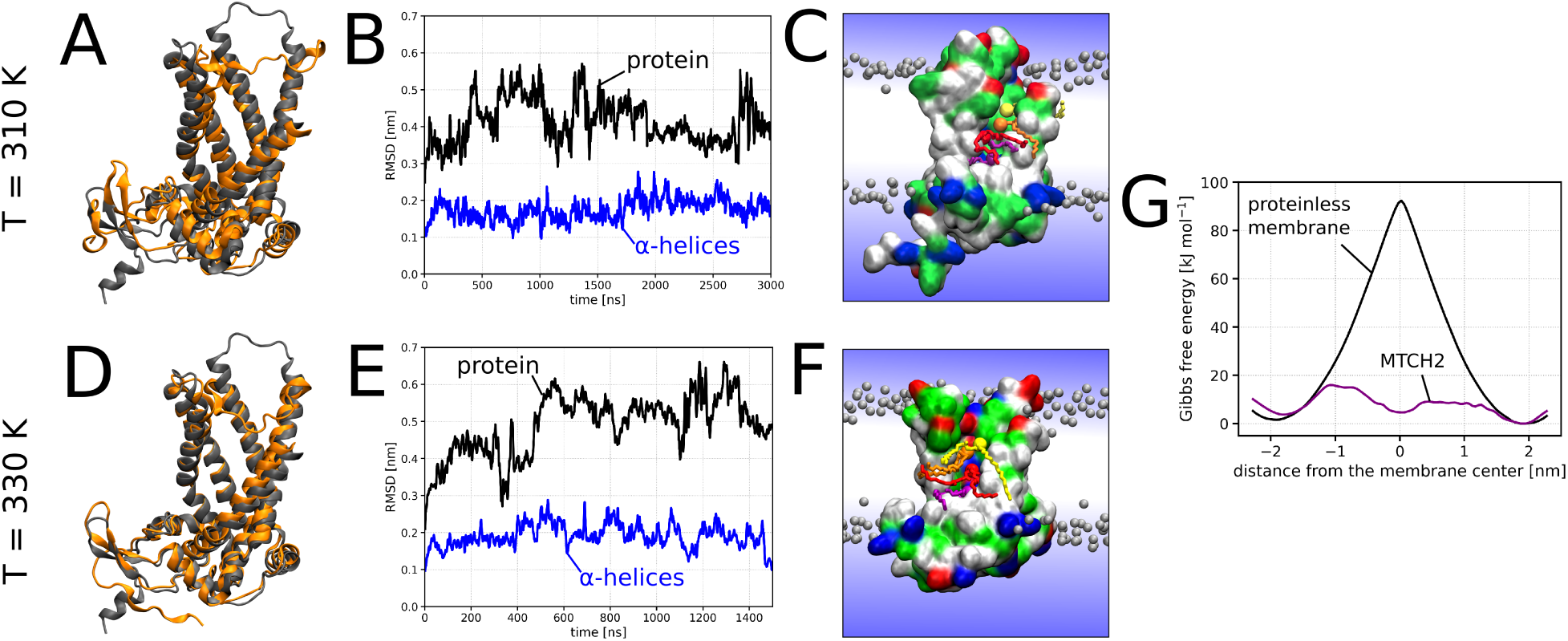
Results of atomistic MD simulations. The upper row (A–C) and the panel G correspond to MTCH2 simulated at 310 K, while the lower row (D–F) corresponds to MTCH2 simulated at 330 K. A and D) Cartoon structure of MTCH2 at the start of the simulation (in gray) overlaid with the structure of MTCH2 at the end of the 3 *μ*s (A) or 1.5 *μ*s (D) atomistic simulation (in orange). Note that the α-helical segments of the protein were quite stable and they did not significantly reorient or reposition during either of the simulations. This suggests that the predicted structure is stable and the protein cavity and the scrambling pathway remain open. B and E) Root-mean-square deviation (RMSD) of the MTCH2 backbone (black) and backbone of α-helices forming the cavity (blue) during the simulation. Note that most of the RMSD calculated for the protein backbone comes from the fluctuations in the loops of MTCH2, as the RMSD for α-helices never reaches 0.3 nm. C and F) Simulation snapshots showing several lipids (colored in yellow, orange, red, and purple) translocating along the scrambling pathway of MTCH2. The protein is shown as a molecular surface colored according to the character of its residues (hydrophilic = green, hydrophobic = white, positively charged = blue, negatively charged = red). Bulk lipids are depicted as gray beads (phosphate group) with hydrophobic tails omitted for clarity. Water is represented only schematically as a blue gradient, omitting the presence of water in the cavity of the protein for visual clarity. G) Free energy profiles of lipid flip-flop in a proteinless POPC membrane (black) and along the MTCH2 scrambling pathway (purple) calculated at T = 310 K. As with Martini simulations, the presence of MTCH2 significantly reduces the flip-flop free energy barrier. Note that the calculation error based on the profile asymmetry is relatively high but still below 5 kJ mol*^−^*^1^. See Figure S3 for detailed view of free energy calculations convergence.

The atomistic simulations also indicated a higher likelihood of lipids and water entering the MTCH2 cavity, as illustrated in Figure 3C. However, we did not observe any full flip-flop event, possibly due to the presence of the “hydrophobic gate” obstructing the lipids from traversing the entire hydrophilic pathway within the limited simulation time. Therefore, we employed umbrella sampling to enhance the scrambling process and calculated the free energy of lipid flip-flop around MTCH2. As shown in Figure 3G, the presence of MTCH2 dramatically decreases the free energy barrier for lipid flip-flop by over 80% compared to the proteinless membrane (from 92 to 16 kJ mol*^−^*^1^), which is in agreement with our Martini simulations. See Figure S7 for representative snapshots from umbrella sampling windows used to calculate the free energy profiles.

Our findings are supported by a recent preprint study, where Li et al.^20^ reported the scrambling activity of MTCH2 both *in vitro* and in unbiased Martini 3 simulations. Using vesicles reconstituted with purified MTCH2 and scramblase assay based on BSA back-extraction of fluorescent lipid reporters^21,22^, they demonstrated that MTCH2 has scramblase activity. Furthermore, via coarse-grained simulations, they identified the same scrambling pathway along MTCH2 as we identified in our study. However, the scrambling rate they reported from Martini 3 simulations, approximately 6 events per *μ*s, is lower than what we observed in our simulations with about 11 scrambling events per *μ*s. This discrepancy is likely due to the different lipid types used in the studies (DOPC in the case of Li et al. versus POPC in our work). Despite this difference, our findings are in line with those of Li et al., and provide more detailed insights into the scrambling pathway and, importantly, the energetics of the scrambling process facilitated by MTCH2.

In our Martini 3 simulations, we have observed that MTCH2 displays a scrambling rate similar to that of the VDAC1 dimer, a recently identified scramblase with a beta-barrel structure^7^. The free energy barriers governing lipid flip-flop in the presence of MTCH2 and a VDAC1 dimer are also closely aligned, with values around 7 kJmol*^−^*^1^ using the Martini 3 model. Both VDAC and MTCH2 are positioned within the outer mitochondrial membrane (OMM), suggesting that MTCH2 could serve as an alternative pathway for lipid scrambling within the OMM. This process is vital for mitochondrial membrane biogenesis and the synthesis of lipids like cardiolipin. However, note that while VDAC requires dimer formation for its scrambling activity, MTCH2 accomplishes lipid scrambling as a monomer, but could be regulated by interaction with other proteins.

In summary, our multiple molecular dynamics simulations and free energy calculations provide evidence for the scrambling activity of MTCH2. Starting from the structure predicted by AlphaFold^11^, we identified a hydrophilic scrambling pathway that shares structural similarities with other known scramblases. Both our coarse-grained and atomistic simulations consistently show the ability of lipids to enter the membrane core along MTCH2. Moreover, our free energy calculations performed with both coarse-grained and atomistic models demonstrate that MTCH2 significantly reduces the free energy barrier for lipid flip-flop. Martini 3 simulations have even resulted in complete lipid flip-flop in both directions along MTCH2, indicating high scrambling activity of this protein. Our findings are in line with the recent experimental evidence for MTCH2 scrambling activity *in vitro* ^20^. The scrambling rate of MTCH2 is similar to that seen with VDAC dimers^7^, which were demonstrated to provide an important mechanism for transporting lipids across the outer membrane of mitochondria. MTCH2 could act redundantly with VDAC, providing complementary scramblase activity. In general, our computational results suggest that insertases such as MTCH2, which possess a hydrophilic path across the membrane, also function as scramblases.

## Acknowledgement

This work was supported by the National Institutes of Health grant NS119779 (AKM), the European Research Council (ERC) under the European Union’s Horizon 2020 research and innovation programme (grant agreement No 101001470) (RV) and the project National Institute of virology and bacteriology (Programme EXCELES, ID Project No. LX22NPO5103) - Funded by the European Union - Next Generation EU (RV). Computational resources were provided by the CESNET, CERIT Scientific Cloud, and IT4 Innovations National Super-computing Center by MEYS CR through the e-INFRA CZ (ID:90254). We acknowledge IT4 Innovations National Supercomputing Center for awarding this project access to the LUMI supercomputer, owned by the EuroHPC Joint Undertaking, hosted by CSC (Finland) and the LUMI consortium through the e-INFRA CZ (ID:90254).

## Author contributions

Conceptualization, A.K.M and R.V.; Methodology, L.B. and R.V.; Software, L.B.; Investigation, L.B.; Resources, L.B. and R.V.; Writing – Original Draft, L.B., A.K.M, and R.V.; Writing – Review & Editing, L.B., A.K.M, and R.V.; Visualization, L.B.; Supervision, A.K.M. and R.V.; Project Administration, A.K.M and R.V.; Funding Acquisition, A.K.M. and R.V.

## Declaration of interests

The authors declare no competing interests.

## Declaration of Generative AI and AI-assisted technologies in the writing process

During the preparation of this work the authors used the ChatGPT tool in order to enhance the clarity, coherence, and overall quality of the writing. After using this tool, the authors reviewed and edited the content as needed and take full responsibility for the content of the publication.

## STAR Methods

## KEY RESOURCES TABLE

| REAGENT or RESOURCE | SOURCE | IDENTIFIER |
| --- | --- | --- |
| Deposited data |  |  |
| Structure of human MTCH2 | AlphaFold Database<sup>12</sup> | Q9Y6C9 |
| Software and algorithms |  |  |
| Gromacs (version 2021.4) | Abraham et al., 2015<sup>23</sup> | https://www.gromacs.org |
| Plumed (version 2.7) | Tribello et al., 2014<sup>24</sup> | https://www.plumed.org |
| CHARMM-GUI | Jo et al., 2008<sup>25</sup> | https://www.charmm-gui.org |
| VMD | Humphrey et al., 1996<sup>26</sup> | https://www.ks.uiuc.edu/Research/vmd |
| martinize2 | Kroon et al., 2023<sup>27</sup> | https://github.com/marrink-lab/vermouth-martinize |
| insane | Wassenaar et al., 2015.<sup>28</sup> | https://github.com/Tsjerk/Insane |
| Grossfield Lab WHAM | Grossfield Lab | http://membrane.urmc.rochester.edu |
| scramblyzer | in-house developed | https://doi.org/10.5281/zenodo.8402719 |

## RESOURCE AVAILABILITY

### Lead contact

Further information and requests should be directed to and will be fulfilled by the lead contact, Robert Vácha.

### Materials availability

This study did not generate new unique reagents.

### Data and code availability

All simulation input files and figure source data are available from Zenodo 10.5281/zen-odo.10159183. In-house developed code for the analysis of the simulations is available from

doi.org/10.5281/zenodo.8402719. All other data such as simulation trajectories are available from the lead contact upon request.

## METHOD DETAILS

### General information

All molecular dynamics (MD) simulations were performed using the simulation package Gromacs version 2021.4^23^ with Plumed plugin version 2.7^24^. All simulation visualizations were prepared using the VMD software^26^. The structure of MTCH2 was obtained from the AlphaFold Protein Structure Database (entry Q9Y6C9) ^11,12^. Prior to all simulations, the obtained structure of MTCH2 was minimized in vacuum using the Amber 99SB-ILDN force field^29^ and the steepest descent algorithm with maximum force tolerance of 100 kJ mol*^−^*^1^ nm*^−^*^1^.

### Martini simulations

We employed two different versions of the Martini force field: Martini version 3^13^ and ElNeDyn force field^30^ based on Martini version 2.2^16–18^ (further called just Martini 2). The minimized atomistic structure of MTCH2 was coarse-grained using martinize2 script^27^ and placed into 1-palmitoyl-2-oleoyl-sn-glycero-3-phosphocholine (POPC) membrane using either insane script (Martini 3)^28^ or CHARMM-GUI (Martini 2)^25^. In Martini 2 simulations, the tertiary structure of MTCH2 was maintained using ElNeDyn, while in Martini 3, we used an elastic network with a force constant of 1000 kJ mol*^−^*^1^ nm*^−^*^2^ to fix the tertiary structure of the protein.

Each Martini system contained one molecule of MTCH2, roughly 920 POPC lipids, about 19,000 water beads and physiological concentration (0.154 mol dm*^−^*^3^) of NaCl ions (with an excess of ions to neutralize the system). The steepest-descent algorithm was used to minimize each system with a maximum force tolerance of 1000 kJ mol*^−^*^1^ nm*^−^*^1^. The equilibration of the systems was performed in five stages, each with different simulation lengths and time steps. The stages were as follows: I) dt = 2 fs, t = 0.5 ns, II) dt = 5 fs, t = 1.25 ns, III) dt = 10 fs, t = 1 ns, IV) dt = 20 fs, t = 30 ns, V) dt = 20 fs, t = 200 ns. During stages I-IV, the Berendsen barostat^31^ was utilized, whereas the Parrinello-Rahman barostat ^32,33^ was used in stage V and in all subsequent simulations. Throughout stages I-IV, a harmonic potential with a force constant of 1000 kJ mol*^−^*^1^ nm*^−^*^2^ was used to restrain all backbone beads of the MTCH2 protein to their initial positions.

We also constructed one proteinless membrane system for each employed Martini force field as a reference. These systems were smaller, composed of roughly 290 POPC lipids, 6000 water beads, and NaCl ions at physiological concentration. The systems were built using insane or CHARMM-GUI (Martini 3 or Martini 2, respectively) and then minimized and equilibrated in the same way as MTCH2 systems, except the stage V of equilibration was shortened to 100 ns. As no protein was present in these systems, no position restraints were applied during the equilibration. The equilibrated structure was then used to perform pulling and subsequently umbrella sampling simulations to obtain the free energy profile of lipid flip-flop in proteinless membrane.

All simulations were performed in the NPT ensemble, with the temperature being maintained at 310 K using stochastic velocity rescaling thermostat^34^ with a coupling constant of 1 ps. Water with ions, membrane, and protein were coupled to three separate thermal baths. Pressure was kept at 1 bar using either the Berendsen^31^ or Parrinello-Rahman barostat ^32,33^ (see above) with a coupling constant of 12 ps. Semi-isotropic pressure coupling was used to independently scale the simulation box in the xy-plane and on the z-axis with a compressibility of 3 *×* 10*^−^*^4^ bar*^−^*^1^. The equations of motion were integrated using the leap-frog algorithm. Non-bonded interactions were cut off at 1.1 nm, and the van der Waals potential was shifted to zero at the cut-off distance. The relative dielectric constant was set to 15 and LINCS^35^ parameters lincs-order and lincs-iter were set to 8 and 2, respectively, to avoid artificial temperature gradients^36^.

Three independent molecular dynamics simulations, each 10 *μ*s long, were run from the equilibrated structure of MTCH2-containing membrane for each of Martini 2 and Martini 3 force fields. Every 10 ns, a simulation snapshot was analyzed to determine the percentage of scrambled lipids over time. A lipid was deemed “scrambled” if it was found in a different membrane leaflet than where it began in the simulation, determined by the position of its phosphate bead in relation to the membrane center. The code for analyzing lipid scrambling is available from doi.org/10.5281/zenodo.8402719.

For the calculation of water and phosphates densities, all three replicas were merged and centered on protein beads based on their root-mean-square deviation. The densities were calculated using VMD Volmap Tool^26^, analyzing a simulation snapshot every 10 ns.

For each of Martini 2 and Martini 3, we calculated the free energy of lipid flip-flop in a) proteinless membrane and b) in membrane containing MTCH2. Umbrella sampling method^37,38^ was used to enhance the sampling of flip-flop events by employing a one-dimensional collective variable (CV). The CV was defined as the oriented distance between the selected lipid phosphate and the local membrane center of mass on the z-axis. The local membrane center of mass was calculated from the positions of lipid beads within a cylinder with a radius of 2.0 nm (for proteinless membrane) or 3.0 nm (for MTCH2 simulations) and its principal axis going through the selected phosphate bead (Gromacs geometry option cylinder).

Initial configurations for umbrella sampling were generated by pulling the selected lipid phosphate bead through the membrane for 1 *μ*s with a pulling rate of 4.6 nm *μ*s*^−^*^1^ (for proteinless membrane) or 4.2 nm *μ*s*^−^*^1^ (for MTCH2 simulations) and initial reference distance of 2.3 nm (for proteinless membrane) or ±2.1 nm (for MTCH2 simulations) using a harmonic potential. For systems with MTCH2, we performed two independent pulling simulations. In one pulling simulation, selected lipid was pulled from the upper to the lower membrane leaflet, while in the other pulling simulation, the lipid was pulled in the opposite direction. 44 (Martini 3 simulations with MTCH2), 64 (Martini 2 simulations with MTCH2), or 67 (simulations of proteinless membrane) umbrella sampling windows distributed along the range of the CV were used. In case of MTCH2 simulations, we generated the windows from pulling simulations performed in each direction. As we observed no hysteresis for Martini 3 simulations with MTCH2, we show both of the calculated free energy profiles. In contrast, Martini 2 simulations with MTCH2 did show a hysteresis (see Figure S6) resulting in a necessity to generate a new set of umbrella sampling windows which initial configurations were obtained from *both* pulling directions. This umbrella sampling simulation was further enhanced by applying Hamiltonian replica exchange^39^ (as implemented in the Plumed plugin^24^) to all the windows with an exchange being attempted every 10,000 simulation steps (200 ps). For Martini 2 simulations with MTCH2, we thus only show one free energy profile in the main text.

See the Tables S1–S3 for the complete list of umbrella sampling windows that were used for the calculations. Each window was simulated for 4 *μ*s (Martini 3 simulations with MTCH2), 3 *μ*s (Martini 2 simulations with MTCH2), or 1 *μ*s (proteinless membrane) with the first 10 ns of simulation time being used for equilibration only. In pulling and umbrella sampling simulations with MTCH2, the selected lipid phosphate was restrained to the backbone bead of F228 using a flat-bottom potential with a reference distance of 2.5 nm in the *xy* plane and a force constant of 500 kJ mol*^−^*^1^ nm*^−^*^2^. The translocating lipid was thus effectively restrained to the scrambling pathway, allowing it to sample the relevant portion of the configuration space.

Free energy profiles were obtained using the weighted histogram analysis method^40,41^ as implemented in Grossfield Lab WHAM program (available from membrane.urmc.rochester.edu).

### Atomistic simulations

For atomistic simulations, we employed the CHARMM36m force field^19^. The MTCH2 structure minimized in vacuum was placed into a POPC membrane using CHARMM-GUI^25^. The system contained one protein, roughly 390 lipid molecules, 37,000 molecules of water, and NaCl ions at a concentration of 0.154 mol dm*^−^*^3^ (with an excess of ions to neutralize the system). Minimization was performed using the steepest-descent algorithm and force tolerance of 1000 kJ mol*^−^*^1^ nm*^−^*^1^. During the minimization, positions restraints were applied to protein backbone and protein sidechains (force constant of 4000 and 2000 kJ mol*^−^*^1^ nm*^−^*^2^, respectively) as well as to the phosphorus atom of each lipid (force constant of 1000 kJ mol*^−^*^1^ nm*^−^*^2^). Dihedral restraints (force constant of 1000 kJ mol*^−^*^1^ rad*^−^*^2^) were further applied to two dihedral angles in all POPC molecules, specifically to dihedrals between C1, C3, C2, and O21 (glycerol carbons and oxygen linking the oleoyl tail to glycerol) and between C28, C29, C210, and C211 (carbons around the double bond of the oleyol tail), fixing them at *−*120^◦^ ± 2.5^◦^ and 0^◦^ ± 0.0^◦^, respectively.

Equilibration was performed in 6 stages of different simulation lengths: I-III) 250 ps (each), IV-V) 1 ns, VI) 5 ns. Stages I-II were performed in the NVT ensemble, while stages III-VI in the NPT ensemble. Stochastic velocity rescaling thermostat^34^ with a coupling constant of 0.5 ps was employed to maintain the temperature of 310 K. Three separate thermal baths were used for water with ions, membrane, and protein. In NPT stages of equilibration, pressure was kept at 1 bar using the Berendsen barostat^31^ with semi-isotropic pressure coupling, coupling constant of 5 ps and compressibility of 4.5 *×* 10*^−^*^5^ bar*^−^*^1^. The equations of motion were integrated using the leap-frog algorithm. In stages I-III, the simulation time step was 1 fs, while in the rest of the equilibration (and the following simulations), the time step was 2 fs. Short-ranged non-bonded interactions were truncated at 1.2 nm, while a force switch was applied starting from 1.0 nm. Electrostatic interactions were treated using Fast Smooth Particle-Mesh Ewald^42^. Bonds with hydrogens were constrained using the LINCS algorithm^35^. Translational velocity removal was applied separately for membrane with protein and for water with ions. Position and dihedral restraints were strong in the initial stages of equilibration and then gradually turned down, see Table S4.

After equilibration, we performed 3 *μ*s long production simulation. In this simulation, the Berendsen thermostat was replaced with Parrinello-Rahman barostat ^32,33^ and no restraints were applied to the system. All the other simulation settings remained the same as in the stage VI of equilibration. RMSD of protein backbone atoms was calculated using the Gromacs rms tool. We also calculated RMSD for α-helices surrounding the cavity which included only backbone atoms of residues 1-31, 74-100, 117-151, 182-207, and 216-247.

We also performed a simulation in which the MTCH2 protein was heated up to further assess its thermodynamic stability. Starting from the minimized structure of the system, we performed a new equilibration with the same simulation settings except the temperature of the protein was increased to 330 K. The same temperature was also applied in the following 1.5 *μ*s long production simulation. The temperature of the membrane and water with ions was kept at the physiological 310 K.

We further calculated free energy of lipid flip-flop in a) a proteinless membrane and b) a membrane containing MTCH2. We used the umbrella sampling method^37,38^ to enhance the sampling of the flip-flop events by employing the same collective variable (CV) as used in the Martini free energy calculations.

For free energy calculations in a proteinless membrane, initial configurations for the umbrella sampling windows were generated by pulling a selected POPC lipid from the upper to the lower membrane leaflet. The pulling rate was set at 9.2 nm *μ*s*^−^*^1^ with an initial reference distance of 2.3 nm and a force constant of 5000 kJ mol*^−^*^1^ nm*^−^*^2^. The pulling was performed over 500 ns. The resulting pulling trajectory was divided into 59 non-uniformly distributed umbrella sampling windows. The spacing ranged from 0.1 nm near the membrane surface to 0.03 nm near the membrane center. A force constant of 1000 kJ mol*^−^*^1^ nm*^−^*^2^ was applied in windows near the membrane surface, while a force constant of 2000 kJ mol*^−^*^1^ nm*^−^*^2^ was used near the membrane center. To enhance sampling further, we employed the Plumed plugin^24^ and applied Hamiltonian replica exchange^39^ to 16 windows in the membrane center (between *−*0.25 and 0.21 nm). Configurations were exchanged every 100,000 integration steps (200 ps). Each umbrella sampling window underwent a simulation of 300 ns, with the first 50 ns used for equilibration only. The complete list of umbrella sampling windows used is presented in Table S5.

For free energy calculations involving MTCH2, multiple pulling simulations were conducted. The first pulling simulation involved pulling a chosen POPC lipid from the upper towards the lower membrane leaflet over approximately 750 ns at a rate of 4.2 nm *μ*s*^−^*^1^, with an initial reference distance of 2.1 nm and a force constant of 5000 kJ mol*^−^*^1^ nm*^−^*^2^. The CV used was the same as in the Martini MTCH2 pulling simulations. The initial configuration for this pulling was the last frame of the 3 *μ*s long production simulation at T = 310 K, described above.

Achieving proper movement of the lipid through the hydrophobic gate proved difficult in the pulling simulation. Therefore, we used the last frame of the first pulling simulation as input for a subsequent pulling simulation, employing a different CV – the *xyz* distance between the chosen lipid’s phosphorus atom and the center of mass of C*_α_* atoms of residues 80-90 of MTCH2. These residues are part of α-helix 3, located opposite to the hydrophobic gate (see Figure 1 B). During this pulling simulation, we increased the distance between the lipid and α-helix 3, forcing the lipid through the hydrophobic gate. The lipid was pulled for 500 ns at a rate of 5.0 nm *μ*s*^−^*^1^ and with a force constant of 5000 kJ mol*^−^*^1^ nm*^−^*^2^, enabling it to cross the hydrophobic gate and reach the lower membrane leaflet.

A third pulling simulation was performed in the opposite direction, starting again from the last frame of the 3 *μ*s long production simulation. We used the same CV as in the second pulling simulation. The CV value was decreased, moving the lipid towards α-helix 3 and through the hydrophobic gate. This pulling was conducted over approximately 500 ns at a rate of 5.0 nm *μ* s*^−^*^1^ and with a force constant of 5000 kJ mol*^−^*^1^ nm*^−^*^2^. Once the lipid reached the cavity of MTCH2, the simulation was terminated.

In all three pulling simulations, as well as in the subsequent umbrella sampling simulation, the phosphorus atom of the pulled lipid was restrained to the backbone bead of F228 of MTCH2 using a flat-bottom potential. The reference distance was set at 2.5 nm in the *xy* plane with a force constant of 500 kJ mol*^−^*^1^ nm*^−^*^2^. This restraint was consistent with the Martini free energy calculations mentioned earlier.

The three pulling trajectories were split into 80 non-uniformly distributed umbrella sampling windows. For umbrella sampling, we used the same CV as in the Martini free energy calculations and the first pulling simulation. In the region of the hydrophobic gate and the C-terminal scrambling pathway (48 umbrella sampling windows), initial configurations were taken from pulling simulations 2 and 3. For the remaining CV range, configurations were taken from the first pulling simulation. Refer to Table S6 for a full list of umbrella sampling windows used. Each window was initially equilibrated for 200 ns. After equilibration, Hamiltonian replica exchange^39^ was applied to 48 umbrella sampling windows around the hydrophobic gate and the C-terminal pathway to further enhance the sampling. Each window then underwent sampling for additional 600 ns, which was utilized to determine the free energy profile.

The approach to obtaining the free energy profiles was consistent with that used in coarsegrained simulations. We utilized the weighted histogram analysis method^40,41^, employing the Grossfield Lab WHAM program (available from membrane.urmc.rochester.edu).

## Supplemental Information

**Table S1:**
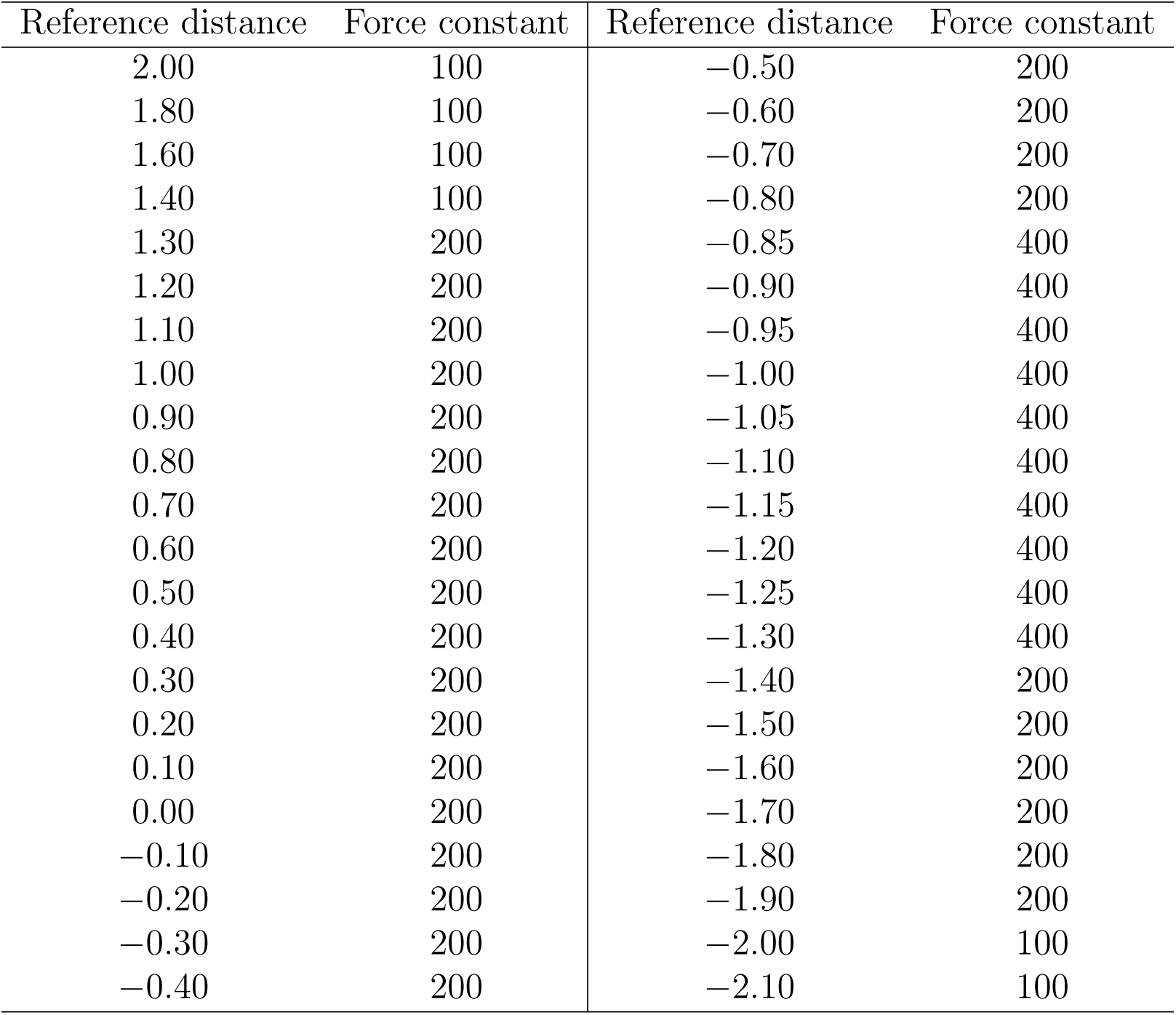
Distribution of umbrella sampling windows along the collective variable with biasing force constants used for Martini 3 simulations of lipid flip-flop through membrane with MTCH2. Reference distances are in nm, force constants in kJ mol*^−^*^1^ nm*^−^*^2^.

| Reference distance | Force constant | Reference distance | Force constant |
| --- | --- | --- | --- |
| 2.00 | 100 | -0.50 | 200 |
| 1.80 | 100 | -0.60 | 200 |
| 1.60 | 100 | -0.70 | 200 |
| 1.40 | 100 | -0.80 | 200 |
| 1.30 | 200 | -0.85 | 400 |
| 1.20 | 200 | -0.90 | 400 |
| 1.10 | 200 | -0.95 | 400 |
| 1.00 | 200 | -1.00 | 400 |
| 0.90 | 200 | -1.05 | 400 |
| 0.80 | 200 | -1.10 | 400 |
| 0.70 | 200 | -1.15 | 400 |
| 0.60 | 200 | -1.20 | 400 |
| 0.50 | 200 | -1.25 | 400 |
| 0.40 | 200 | -1.30 | 400 |
| 0.30 | 200 | -1.40 | 200 |
| 0.20 | 200 | -1.50 | 200 |
| 0.10 | 200 | -1.60 | 200 |
| 0.00 | 200 | -1.70 | 200 |
| -0.10 | 200 | -1.80 | 200 |
| -0.20 | 200 | -1.90 | 200 |
| -0.30 | 200 | -2.00 | 100 |
| -0.40 | 200 | -2.10 | 100 |

**Table S2:**
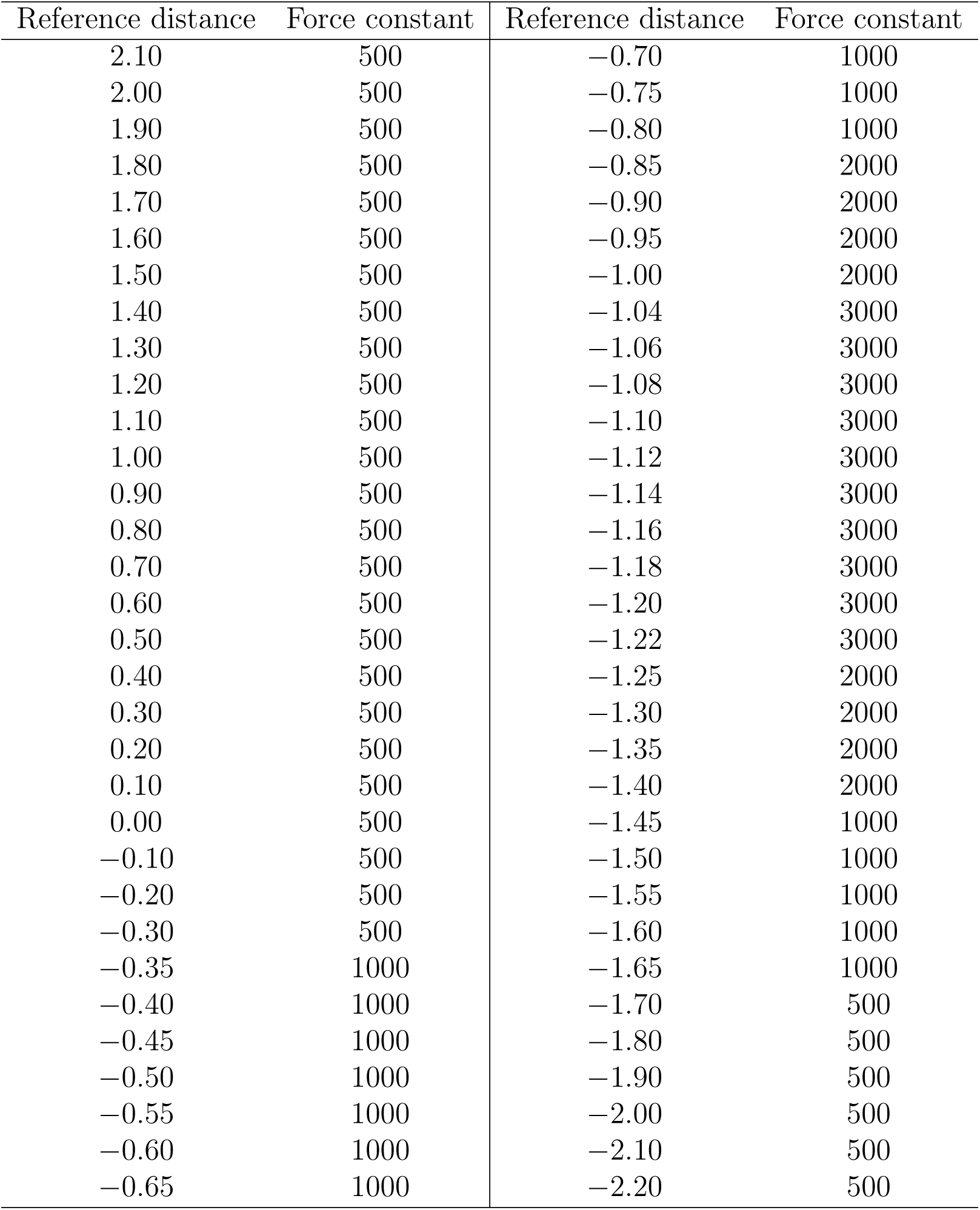
Distribution of umbrella sampling windows along the collective variable with biasing force constants used for Martini 2 simulations of lipid flip-flip through membrane with MTCH2. Reference distances are in nm, force constants in kJ mol*^−^*^1^ nm*^−^*^2^.

**Table S3:**
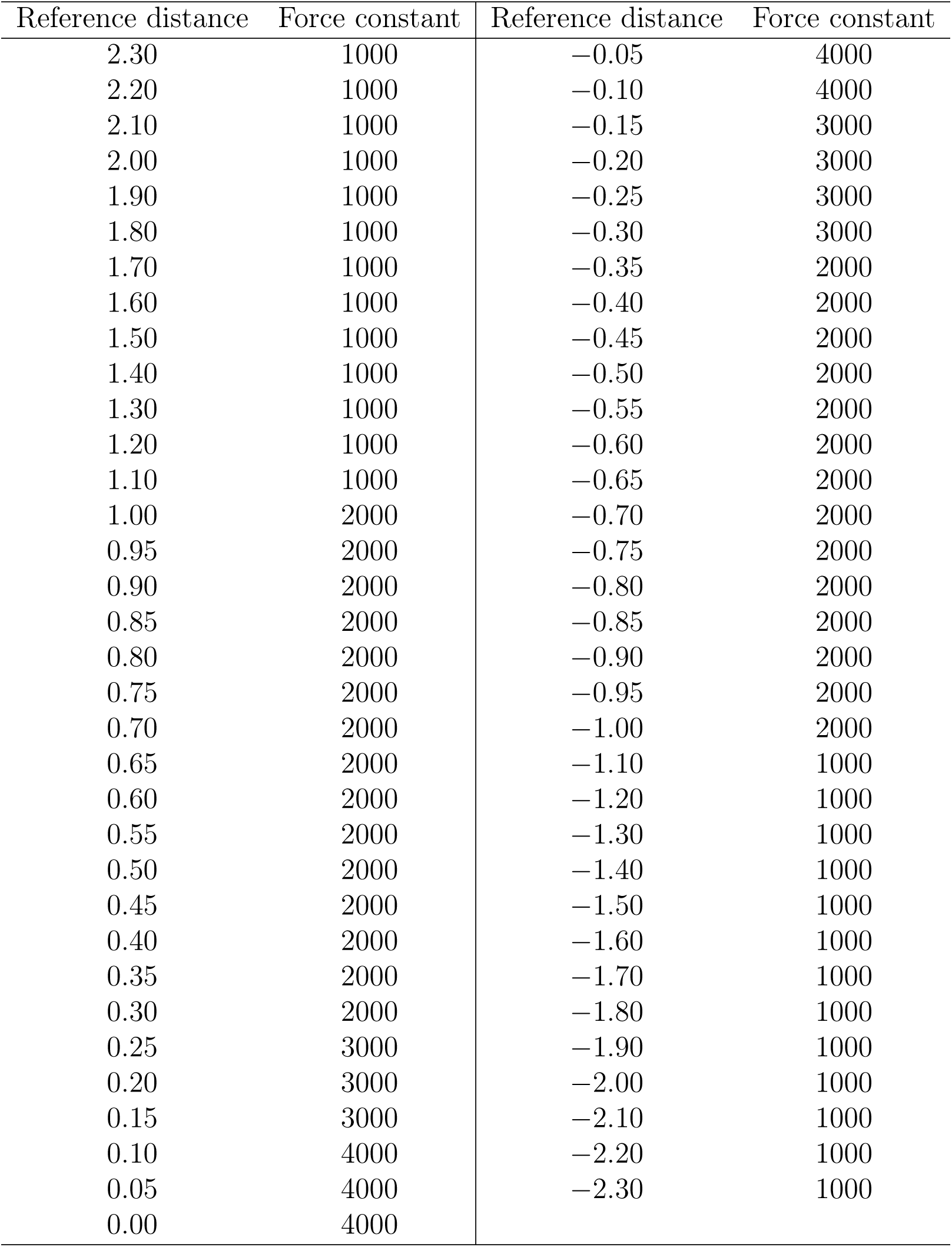
Distribution of umbrella sampling windows along the collective variable with biasing force constants used for Martini 2 and 3 simulations of lipid flip-flop through proteinless membrane. Reference distances are in nm, force constants in kJ mol*^−^*^1^ nm*^−^*^2^.

| Reference distance | Force constant | Reference distance | Force constant |
| --- | --- | --- | --- |
| 2.30 | 1000 | -0.05 | 4000 |
| 2.20 | 1000 | -0.10 | 4000 |
| 2.10 | 1000 | -0.15 | 3000 |
| 2.00 | 1000 | -0.20 | 3000 |
| 1.90 | 1000 | -0.25 | 3000 |
| 1.80 | 1000 | -0.30 | 3000 |
| 1.70 | 1000 | -0.35 | 2000 |
| 1.60 | 1000 | -0.40 | 2000 |
| 1.50 | 1000 | -0.45 | 2000 |
| 1.40 | 1000 | -0.50 | 2000 |
| 1.30 | 1000 | -0.55 | 2000 |
| 1.20 | 1000 | -0.60 | 2000 |
| 1.10 | 1000 | -0.65 | 2000 |
| 1.00 | 2000 | -0.70 | 2000 |
| 0.95 | 2000 | -0.75 | 2000 |
| 0.90 | 2000 | -0.80 | 2000 |
| 0.85 | 2000 | -0.85 | 2000 |
| 0.80 | 2000 | -0.90 | 2000 |
| 0.75 | 2000 | -0.95 | 2000 |
| 0.70 | 2000 | -1.00 | 2000 |
| 0.65 | 2000 | -1.10 | 1000 |
| 0.60 | 2000 | -1.20 | 1000 |
| 0.55 | 2000 | -1.30 | 1000 |
| 0.50 | 2000 | -1.40 | 1000 |
| 0.45 | 2000 | -1.50 | 1000 |
| 0.40 | 2000 | -1.60 | 1000 |
| 0.35 | 2000 | -1.70 | 1000 |
| 0.30 | 2000 | -1.80 | 1000 |
| 0.25 | 3000 | -1.90 | 1000 |
| 0.20 | 3000 | -2.00 | 1000 |
| 0.15 | 3000 | -2.10 | 1000 |
| 0.10 | 4000 | -2.20 | 1000 |
| 0.05 | 4000 | -2.30 | 1000 |
| 0.00 | 4000 |  |  |

**Table S4:** Position and dihedral restraints applied during the individual stages of atomistic equilibration. All values are in kJ mol*^−^*^1^ nm*^−^*^2^ or kJ mol*^−^*^1^ rad*^−^*^2^ (for lipid dihedrals).

|  | protein backbone | protein sidechains | lipid phosphorus | lipid dihedrals |
| --- | --- | --- | --- | --- |
| I | 4000 | 2000 | 1000 | 1000 |
| II | 2000 | 1000 | 400 | 400 |
| III | 1000 | 500 | 400 | 200 |
| IV | 500 | 200 | 200 | 200 |
| V | 200 | 50 | 40 | 100 |
| VI | 50 | 0 | 0 | 0 |

**Table S5:** Distribution of umbrella sampling windows along the collective variable with biasing force constants used for atomistic simulations of lipid flip-flop through proteinless membrane. *^R^* marks windows where Hamiltonian replica exchange was applied. Reference distances are in nm, force constants in kJ mol*^−^*^1^ nm*^−^*^2^.

| Reference distance | Force constant | Reference distance | Force constant |
| --- | --- | --- | --- |
| 2.30 | 1000 | -0.03 | 2000<sup>R</sup> |
| 2.20 | 1000 | -0.06 | 2000<sup>R</sup> |
| 2.10 | 1000 | -0.09 | 2000<sup>R</sup> |
| 2.00 | 1000 | -0.12 | 2000<sup>R</sup> |
| 1.90 | 1000 | -0.15 | 2000<sup>R</sup> |
| 1.80 | 1000 | -0.18 | 2000<sup>R</sup> |
| 1.70 | 1000 | -0.21 | 2000<sup>R</sup> |
| 1.60 | 1000 | -0.25 | 2000<sup>R</sup> |
| 1.50 | 1000 | -0.30 | 1000 |
| 1.40 | 1000 | -0.40 | 1000 |
| 1.30 | 1000 | -0.50 | 1000 |
| 1.20 | 1000 | -0.60 | 1000 |
| 1.10 | 1000 | -0.70 | 1000 |
| 1.00 | 1000 | -0.80 | 1000 |
| 0.90 | 1000 | -0.90 | 1000 |
| 0.80 | 1000 | -1.00 | 1000 |
| 0.70 | 1000 | -1.10 | 1000 |
| 0.60 | 1000 | -1.20 | 1000 |
| 0.50 | 1000 | -1.30 | 1000 |
| 0.40 | 1000 | -1.40 | 1000 |
| 0.30 | 1000 | -1.50 | 1000 |
| 0.25 | 2000 | -1.60 | 1000 |
| 0.21 | 2000<sup>R</sup> | -1.70 | 1000 |
| 0.18 | 2000<sup>R</sup> | -1.80 | 1000 |
| 0.15 | 2000<sup>R</sup> | -1.90 | 1000 |
| 0.12 | 2000<sup>R</sup> | -2.00 | 1000 |
| 0.09 | 2000<sup>R</sup> | -2.10 | 1000 |
| 0.06 | 2000<sup>R</sup> | -2.20 | 1000 |
| 0.03 | 2000<sup>R</sup> | -2.30 | 1000 |
| 0.00 | 2000<sup>R</sup> |  |  |

**Table S6:** Distribution of umbrella sampling windows along the collective variable used for atomistic simulations of lipid flip-flop through membrane with MTCH2. “Origin” corresponds to the pulling simulation from which the initial configuration for the given umbrella sampling window was taken. *^R^* marks windows where Hamiltonian replica exchange was applied. Reference distances are in nm, force constants in kJ mol*^−^*^1^ nm*^−^*^2^.

| Reference distance | Force constant | Origin | Reference distance | Force constant | Origin |
| --- | --- | --- | --- | --- | --- |
| 2.20 | 1000 | pull 1 | -0.90 | 1000 | pull 2<sup>R</sup> |
| 2.10 | 1000 | pull 1 | -0.92 | 3000 | pull 3<sup>R</sup> |
| 2.00 | 1000 | pull 1 | -0.94 | 3000 | pull 2<sup>R</sup> |
| 1.90 | 1000 | pull 1 | -0.96 | 3000 | pull 3<sup>R</sup> |
| 1.80 | 1000 | pull 1 | -0.98 | 3000 | pull 2<sup>R</sup> |
| 1.70 | 1000 | pull 1 | -1.00 | 3000 | pull 3<sup>R</sup> |
| 1.60 | 1000 | pull 1 | -1.02 | 3000 | pull 2<sup>R</sup> |
| 1.50 | 1000 | pull 1 | -1.04 | 3000 | pull 3<sup>R</sup> |
| 1.40 | 1000 | pull 1 | -1.06 | 3000 | pull 2<sup>R</sup> |
| 1.30 | 1000 | pull 1 | -1.08 | 3000 | pull 3<sup>R</sup> |
| 1.20 | 1000 | pull 1 | -1.10 | 3000 | pull 2<sup>R</sup> |
| 1.10 | 1000 | pull 1 | -1.12 | 3000 | pull 3<sup>R</sup> |
| 1.00 | 1000 | pull 1 | -1.14 | 3000 | pull 2<sup>R</sup> |
| 0.90 | 1000 | pull 1 | -1.16 | 3000 | pull 3<sup>R</sup> |
| 0.80 | 1000 | pull 1 | -1.18 | 3000 | pull 2<sup>R</sup> |
| 0.70 | 1000 | pull 1 | -1.20 | 3000 | pull 3<sup>R</sup> |
| 0.60 | 1000 | pull 1 | -1.22 | 3000 | pull 2<sup>R</sup> |
| 0.50 | 1000 | pull 1 | -1.24 | 3000 | pull 3<sup>R</sup> |
| 0.40 | 1000 | pull 1 | -1.26 | 3000 | pull 2<sup>R</sup> |
| 0.30 | 1000 | pull 1 | -1.28 | 3000 | pull 3<sup>R</sup> |
| 0.20 | 1000 | pull 1 | -1.30 | 3000 | pull 2<sup>R</sup> |
| 0.10 | 1000 | pull 1 | -1.32 | 3000 | pull 3<sup>R</sup> |
| 0.00 | 1000 | pull 1 | -1.34 | 3000 | pull 2<sup>R</sup> |
| -0.05 | 1000 | pull 1 | -1.36 | 3000 | pull 3<sup>R</sup> |
| -0.10 | 1000 | pull 1 | -1.38 | 3000 | pull 2<sup>R</sup> |
| -0.15 | 1000 | pull 1 | -1.40 | 1000 | pull 3<sup>R</sup> |
| -0.20 | 1000 | pull 1 | -1.45 | 1000 | pull 2<sup>R</sup> |
| -0.25 | 1000 | pull 1 | -1.50 | 1000 | pull 3<sup>R</sup> |
| -0.30 | 1000 | pull 1 | -1.55 | 1000 | pull 2<sup>R</sup> |
| -0.35 | 1000 | pull 1 | -1.60 | 1000 | pull 3<sup>R</sup> |
| -0.40 | 1000 | pull 1 | -1.65 | 1000 | pull 2<sup>R</sup> |
| -0.45 | 1000 | pull 1 | -1.70 | 1000 | pull 3<sup>R</sup> |
| -0.50 | 1000 | pull 2<sup>R</sup> | -1.75 | 1000 | pull 2<sup>R</sup> |
| -0.55 | 1000 | pull 3<sup>R</sup> | -1.80 | 1000 | pull 3<sup>R</sup> |
| -0.60 | 1000 | pull 2<sup>R</sup> | -1.85 | 1000 | pull 2<sup>R</sup> |
| -0.65 | 1000 | pull 3<sup>R</sup> | -1.90 | 1000 | pull 3<sup>R</sup> |
| -0.70 | 1000 | pull 2<sup>R</sup> | -1.95 | 1000 | pull 2<sup>R</sup> |
| -0.75 | 1000 | pull 3<sup>R</sup> | -2.00 | 500 | pull 3<sup>R</sup> |
| -0.80 | 1000 | pull 2<sup>R</sup> | -2.10 | 500 | pull 2<sup>R</sup> |
| -0.85 | 1000 | pull 3<sup>R</sup> | -2.20 | 500 | pull 3<sup>R</sup> |

**Figure S1:**
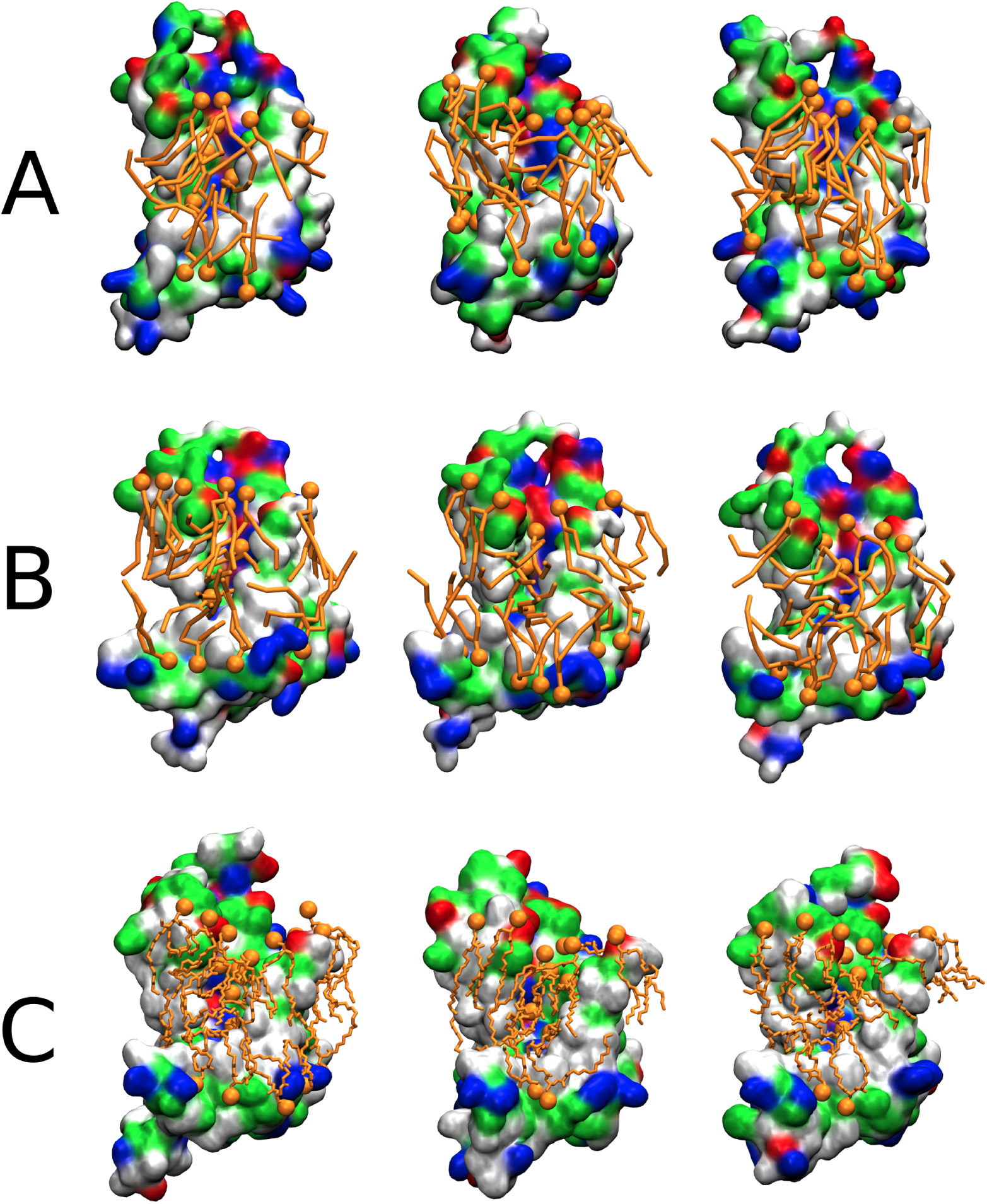
Simulation snapshots obtained from unbiased Martini 3 (A), Martini 2 (B), and atomistic (C) simulations showing insertion of lipid headgroups into the hydrophilic cavity of MTCH2 and the disorder in the lipid tails. Only lipids in proximity to the cavity are shown. Lipid headgroups are depicted as orange beads while lipid tails as orange tubes. MTCH2 is represented as a molecular surface colored according to the character of its residues (hydrophilic = green, hydrophobic = white, positively charged = blue, negatively charged = red).

**Figure S2:**
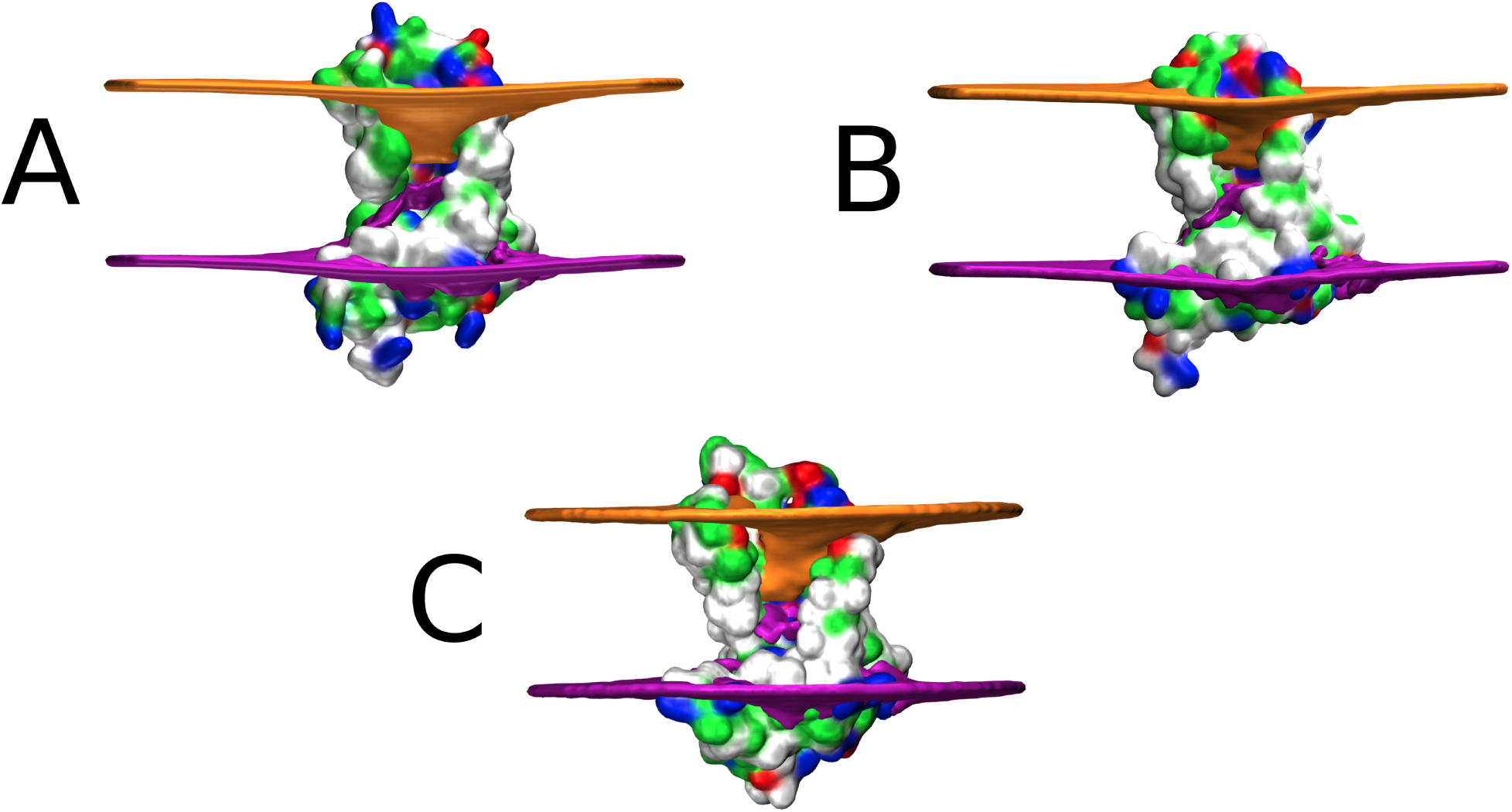
Membrane thickness in the vicinity of MTCH2 calculated from unbiased Martini 3 (A), Martini 2 (B), and atomistic (C) simulations. The orange and purple surfaces represent the average positions of lipid phosphate beads (or phosphorus atoms, in case of atomistic simulation) in the upper and lower leaflet, respectively. The membrane is dramatically thinned in the cavity of MTCH2 in all three force fields.

**Figure S3:**
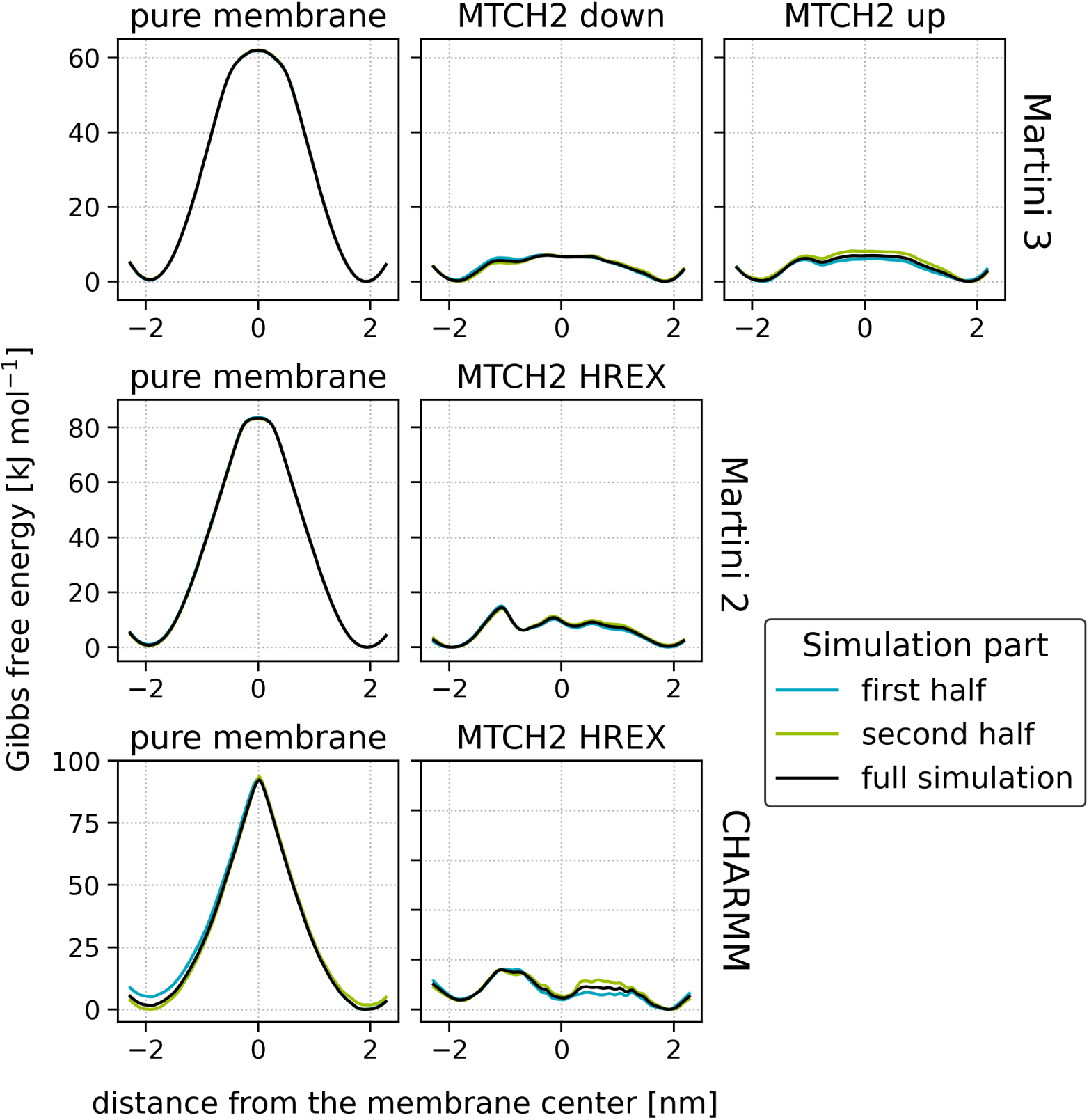
Convergence of free energy calculations captured by calculating additional free energy profiles from the first (azure) and the second (lime) halves of the production phase of umbrella sampling. “MTCH2 down” and “MTCH2 up” correspond to umbrella sampling simulations in which MTCH2 was present, and the initial configurations for the umbrella sampling windows were obtained from steered molecular dynamics, during which the selected lipid phosphate was pulled from the upper to the lower leaflet and from the lower to the upper leaflet, respectively. “MTCH2 HREX” corresponds to simulation in which umbrella sampling was enhanced by Hamiltonian replica exchange. For more details, refer to the Methods section. Note that an additional estimate of the error can be obtained from the asymmetry of the free energy minima located on each side of the profile.

**Figure S4:**
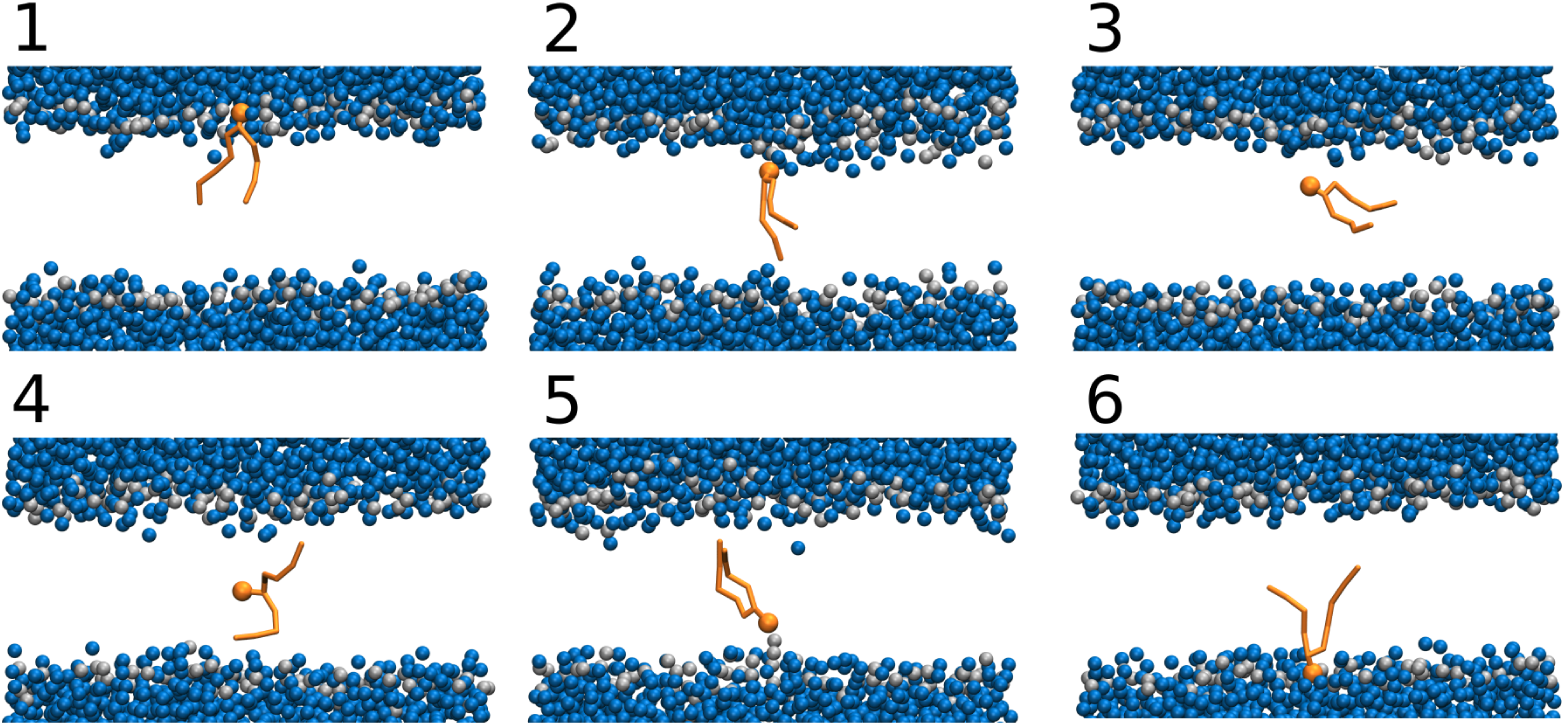
Representative simulation snapshots of a selected lipid translocating through a proteinless membrane captured from Martini 3 umbrella sampling windows. The snapshots are centered on the phosphate of the translocating lipid. The translocating lipid is shown in orange, while only phosphate groups are shown for the other lipids, represented by gray beads. Only water beads close to the membrane are shown, and these are depicted as blue beads.

**Figure S5:**
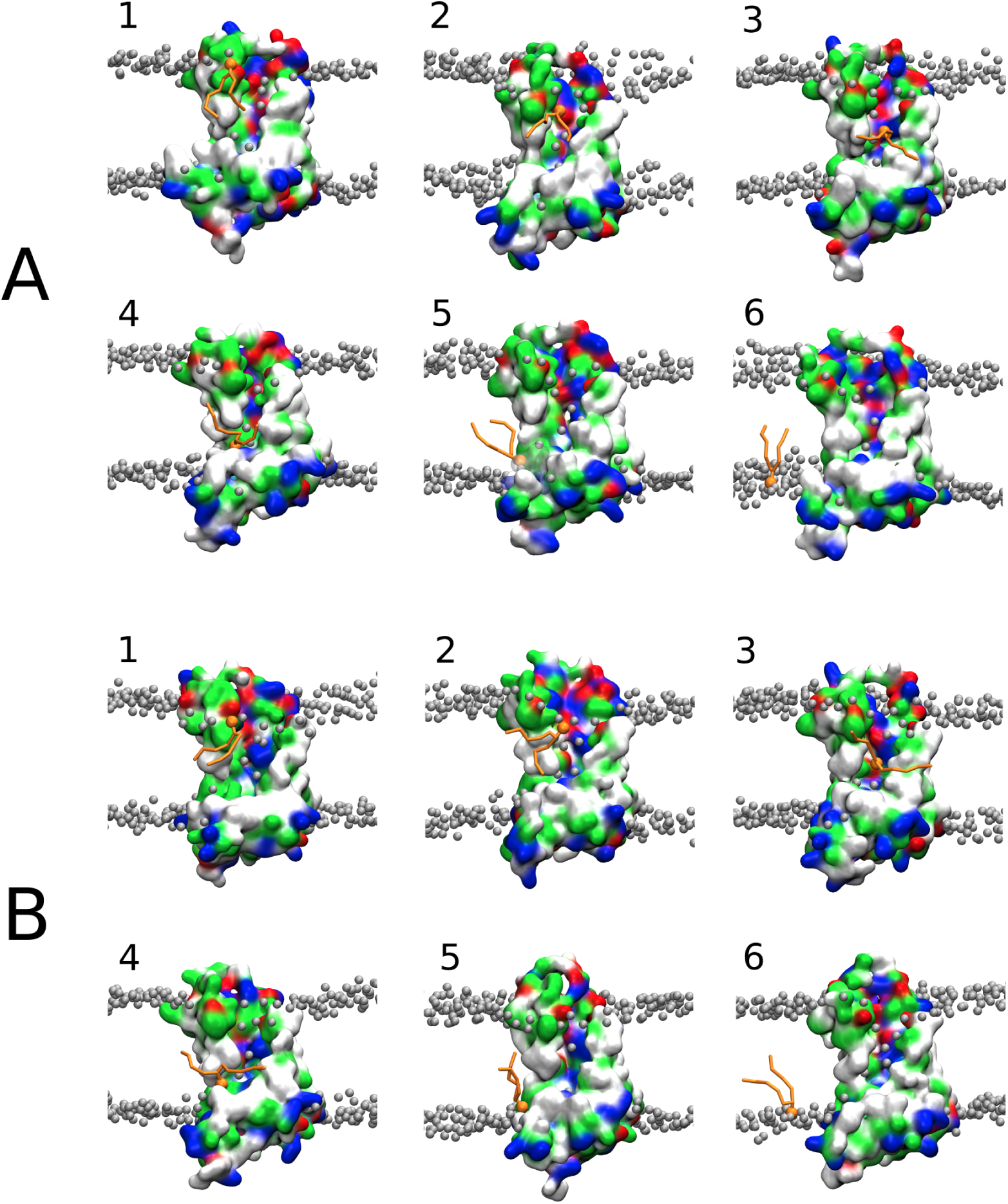
Representative simulation snapshots captured from Martini 3 umbrella sampling windows. The initial configurations for the umbrella sampling windows were obtained from steered molecular dynamics (MD) simulations which involved the pulling of a selected lipid phosphate either from the upper leaflet to the lower leaflet (A) or from the lower leaflet to the upper leaflet (B). MTCH2 is depicted as a molecular surface colored according to the character of its residues (hydrophilic = green, hydrophobic = white, positively charged = blue, negatively charged = red). The pulled lipid is shown in orange, while other lipids are only partially represented with their phosphate groups illustrated as gray beads. Water and ions are not shown.

**Figure S6:**
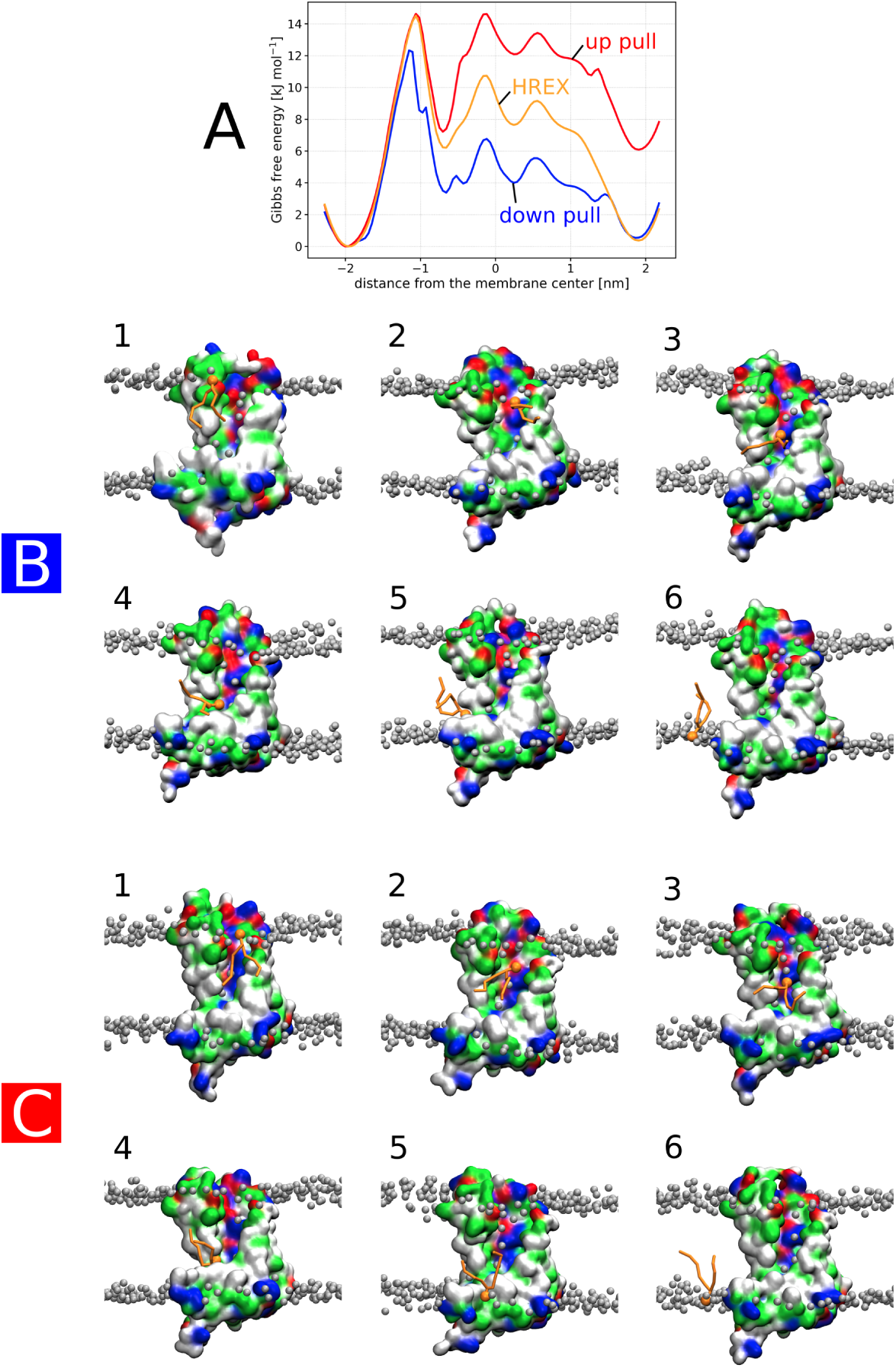
A) Comparison of free energy profiles calculated using umbrella sampling windows with initial configurations obtained from steered molecular dynamics simulations, where the selected lipid phosphate was either pulled from the upper to the lower leaflet (down pull, blue profile) or from the lower to the upper leaflet (up pull, red profile). The HREX, orange profile was calculated using umbrella sampling windows from both pulling directions (by alternating the origin of initial configurations) while employing Hamiltonian replica exchange. Note the hysteresis between the red and blue profiles, which prompted the use of Hamiltonian replica exchange. B–C) Representative simulation snapshots captured from Martini 2 umbrella sampling windows. B corresponds to the set of windows originating from the “down pull”, while C corresponds to windows originating from the “up pull”. “HREX” windows are not shown. The color scheme is the same as in Figure S5. The difference between the “down pull” and “up pull” profiles likely originates from the lipid using a different scrambling pathway in the C-terminal part of the protein when pulled from the lower to the upper leaflet (compare snapshots B5 and C5). Note that this alternative pathway is likely an artifact of pulling, as it was not observed in the windows with applied Hamiltonian replica exchange after initial equilibration.

**Figure S7:**
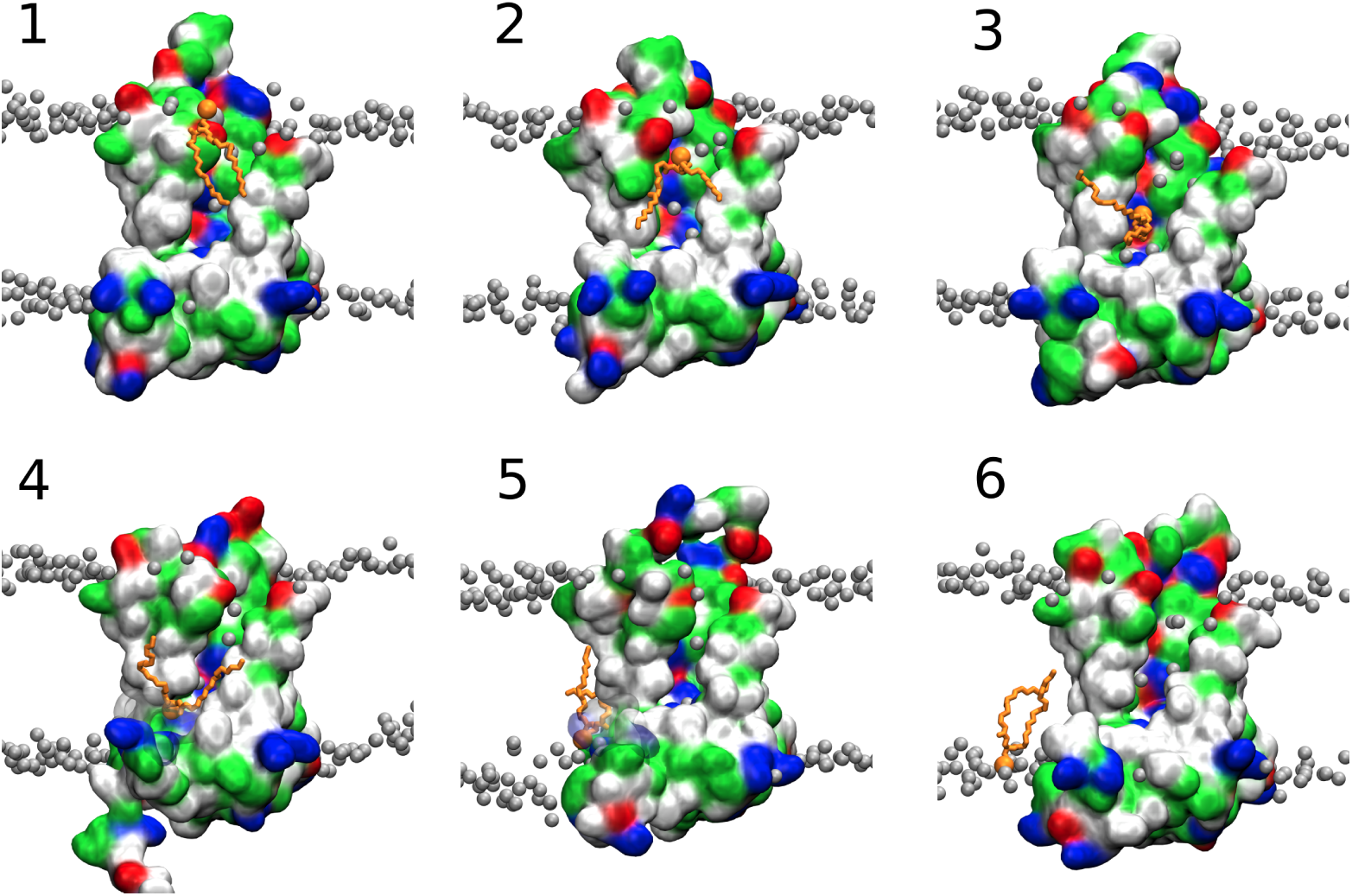
Representative simulation snapshots of a selected lipid translocating through a membrane containing MTCH2 captured from atomistic umbrella sampling windows. MTCH2 is depicted as a molecular surface colored according to the character of its residues (hydrophilic = green, hydrophobic = white, positively charged = blue, negatively charged = red). The pulled lipid is shown in orange, while other lipids are only partially represented with their phosphate groups illustrated as gray beads. Water and ions are not shown.

## Notes

https://doi.org/10.5281/zenodo.10159183

### Competing Interest Statement

The authors have declared no competing interest.

### Summary of Updates

Added free energy profiles using all-atom simulations

## References

(1) Pomorski, T.; Menon, A. K. Lipid flippases and their biological functions. Cellular and Molecular Life Sciences CMLS 2006, 63, 2908–2921.

(2) Sakuragi, T.; Nagata, S. Regulation of phospholipid distribution in the lipid bilayer by flippases and scramblases. Nature Reviews Molecular Cell Biology 2023, 24, 576–596.

(3) Kalienkova, V.; Clerico Mosina, V.; Paulino, C. The Groovy TMEM16 Family: Molecular Mechanisms of Lipid Scrambling and Ion Conduction. Journal of Molecular Biology 2021, 433, 166941.

(4) Khelashvili, G.; Menon, A. K. Phospholipid Scrambling by G Protein–Coupled Receptors. Annual Review of Biophysics 2022, 51 .

(5) Okawa, F.; Hama, Y.; Zhang, S.; Morishita, H.; Yamamoto, H.; Levine, T. P.; Mizushima, N. Evolution and insights into the structure and function of the DedA superfamily containing TMEM41B and VMP1. Journal of Cell Science 2021, 134, jcs255877.

(6) Wang, Y.; Menon, A. K.; Maki, Y.; Liu, Y.-S.; Iwasaki, Y.; Fujita, M.; Guerrero, P. A.; Silva, D. V.; Seeberger, P. H.; Murakami, Y. et al. Genome-wide CRISPR screen reveals CLPTM1L as a lipid scramblase required for efficient glycosylphosphatidylinositol biosynthesis. Proceedings of the National Academy of Sciences 2022, 119, e2115083119.

(7) Jahn, H.; Bartoš, L.; Dearden, G. I.; Dittman, J. S.; Holthuis, J. C. M.; Vácha, R.; Menon, A. K. Phospholipids are imported into mitochondria by VDAC, a dimeric beta barrel scramblase. bioRxiv 2023,

(8) Kizmaz, B.; Herrmann, J. M. Membrane insertases at a glance. Journal of Cell Science 2023, 136, jcs261219.

(9) Wu, X.; Rapoport, T. A. Translocation of Proteins through a Distorted Lipid Bilayer. Trends in Cell Biology 2021, 31, 473–484.

(10) Guna, A.; Stevens, T. A.; Inglis, A. J.; Replogle, J. M.; Esantsi, T. K.; Muthukumar, G.; Shaffer, K. C. L.; Wang, M. L.; Pogson, A. N.; Jones, J. J. et al. MTCH2 is a mitochondrial outer membrane protein insertase. Science 2022, 378, 317–322.

(11) Jumper, J.; Evans, R.; Pritzel, A.; Green, T.; Figurnov, M.; Ronneberger, O.; Tunyasuvunakool, K.; Bates, R.; Žídek, A.; Potapenko, A., et al. Highly accurate protein structure prediction with AlphaFold. Nature 2021, 596, 583–589.

(12) Varadi, M.; Anyango, S.; Deshpande, M.; Nair, S.; Natassia, C.; Yordanova, G.; Yuan, D.; Stroe, O.; Wood, G.; Laydon, A. et al. AlphaFold Protein Structure Database: massively expanding the structural coverage of protein-sequence space with high-accuracy models. Nucleic Acids Research 2022, 50, D439–D444.

(13) Souza, P. C. T.; Alessandri, R.; Barnoud, J.; Thallmair, S.; Faustino, I.; Grünewald, F.; Patmanidis, I.; Abdizadeh, H.; Bruininks, B. M. H.; Wassenaar, T. A. et al. Martini 3: a general purpose force field for coarse-grained molecular dynamics. Nature Methods 2021, 18, 382–388.

(14) Brunner, J. D.; Lim, N. K.; Schenck, S.; Duerst, A.; Dutzler, R. X-ray structure of a calcium-activated TMEM16 lipid scramblase. Nature 2014, 516, 207–212.

(15) Morra, G.; Razavi, A. M.; Pandey, K.; Weinstein, H.; Menon, A. K.; Khelashvili, G. Mechanisms of Lipid Scrambling by the G Protein-Coupled Receptor Opsin. Structure 2018, 26, 356–367.e3.

(16) Marrink, S. J.; Risselada, H. J.; Yefimov, S.; Tieleman, D. P.; de Vries, A. H. The MARTINI Force Field: Coarse Grained Model for Biomolecular Simulations. The Journal of Physical Chemistry B 2007, 111, 7812–7824.

(17) Monticelli, L.; Kandasamy, S. K.; Periole, X.; Larson, R. G.; Tieleman, D. P.; Marrink, S.-J. The MARTINI Coarse-Grained Force Field: Extension to Proteins. Journal of Chemical Theory and Computation 2008, 4, 819–834.

(18) de Jong, D. H.; Singh, G.; Bennett, W. F. D.; Arnarez, C.; Wassenaar, T. A.; Schäfer, L. V.; Periole, X.; Tieleman, D. P.; Marrink, S. J. Improved Parameters for the Martini Coarse-Grained Protein Force Field. Journal of Chemical Theory and Computation 2013, 9, 687–697.

(19) Huang, J.; Rauscher, S.; Nawrocki, G.; Ran, T.; Feig, M.; de Groot, B. L.; Grubmüller, H.; MacKerell, A. D. CHARMM36m: an improved force field for folded and intrinsically disordered proteins. Nature Methods 2017, 14, 71–73.

(20) Li, D.; Rocha-Roa, C.; Schilling, M. A.; Reinisch, K. M.; Vanni, S. Lipid scrambling is a general feature of protein insertases. bioRxiv 2023,

(21) Kubelt, J.; Menon, A. K.; Müller, P.; Herrmann, A. Transbilayer Movement of Fluorescent Phospholipid Analogues in the Cytoplasmic Membrane of *Escherichia coli*. Biochemistry 2002, 41, 5605–5612.

(22) Chang, Q.-l.; Gummadi, S. N.; Menon, A. K. Chemical Modification Identifies Two Populations of Glycerophospholipid Flippase in Rat Liver ER. Biochemistry 2004, 43, 10710–10718.

(23) Abraham, M. J.; Murtola, T.; Schulz, R.; Páll, S.; Smith, J. C.; Hess, B.; Lindahl, E. GROMACS: High performance molecular simulations through multi-level parallelism from laptops to supercomputers. SoftwareX 2015, *1-2*, 19–25.

(24) Tribello, G. A.; Bonomi, M.; Branduardi, D.; Camilloni, C.; Bussi, G. PLUMED 2: New feathers for an old bird. Computer Physics Communications 2014, 185, 604–613.

(25) Jo, S.; Kim, T.; Iyer, V. G.; Im, W. CHARMM-GUI: A web-based graphical user interface for CHARMM. Journal of Computational Chemistry 2008, 29, 1859–1865.

(26) Humphrey, W.; Dalke, A.; Schulten, K. VMD: Visual molecular dynamics. Journal of Molecular Graphics 1996, 14, 33–38.

(27) Kroon, P. C.; Grunewald, F.; Barnoud, J.; Tilburg, M. v.; Souza, P. C. T.; Wassenaar, T. A.; Marrink, S. J. Martinize2 and Vermouth: Unified Framework for Topology Generation. eLife 2023, 12, RP90627.

(28) Wassenaar, T. A.; Ingólfsson, H. I.; Böckmann, R. A.; Tieleman, D. P.; Marrink, S. J. Computational Lipidomics with *insane* : A Versatile Tool for Generating Custom Membranes for Molecular Simulations. Journal of Chemical Theory and Computation 2015, 11, 2144–2155.

(29) Lindorff-Larsen, K.; Piana, S.; Palmo, K.; Maragakis, P.; Klepeis, J. L.; Dror, R. O.; Shaw, D. E. Improved side-chain torsion potentials for the Amber ff99SB protein force field: Improved Protein Side-Chain Potentials. Proteins: Structure, Function, and Bioinformatics 2010, 78, 1950–1958.

(30) Periole, X.; Cavalli, M.; Marrink, S.-J.; Ceruso, M. A. Combining an Elastic Network With a Coarse-Grained Molecular Force Field: Structure, Dynamics, and Intermolecular Recognition. Journal of Chemical Theory and Computation 2009, 5, 2531–2543.

(31) Berendsen, H. J. C.; Postma, J. P. M.; van Gunsteren, W. F.; DiNola, A.; Haak, J. R. Molecular dynamics with coupling to an external bath. The Journal of Chemical Physics 1984, 81, 3684–3690.

(32) Parrinello, M.; Rahman, A. Crystal Structure and Pair Potentials: A Molecular-Dynamics Study. Physical Review Letters 1980, 45, 1196–1199.

(33) Parrinello, M.; Rahman, A. Polymorphic transitions in single crystals: A new molecular dynamics method. Journal of Applied Physics 1981, 52, 7182–7190.

(34) Bussi, G.; Donadio, D.; Parrinello, M. Canonical sampling through velocity rescaling. The Journal of Chemical Physics 2007, 126, 014101.

(35) Hess, B.; Bekker, H.; Berendsen, H. J. C.; Fraaije, J. G. E. M. LINCS: A linear constraint solver for molecular simulations. Journal of Computational Chemistry 1997, 18, 1463–1472.

(36) Thallmair, S.; Javanainen, M.; Fábián, B.; Martinez-Seara, H.; Marrink, S. J. Non-converged Constraints Cause Artificial Temperature Gradients in Lipid Bilayer Simulations. The Journal of Physical Chemistry B 2021, 125, 9537–9546.

(37) Torrie, G.; Valleau, J. Monte Carlo free energy estimates using non-Boltzmann sampling: Application to the sub-critical Lennard-Jones fluid. Chemical Physics Letters 1974, 28, 578–581.

(38) Torrie, G.; Valleau, J. Nonphysical sampling distributions in Monte Carlo free-energy estimation: Umbrella sampling. Journal of Computational Physics 1977, 23, 187–199.

(39) Fukunishi, H.; Watanabe, O.; Takada, S. On the Hamiltonian replica exchange method for efficient sampling of biomolecular systems: Application to protein structure prediction. The Journal of Chemical Physics 2002, 116, 9058–9067.

(40) Kumar, S.; Rosenberg, J. M.; Bouzida, D.; Swendsen, R. H.; Kollman, P. A. The weighted histogram analysis method for free-energy calculations on biomolecules. I. The method. Journal of Computational Chemistry 1992, 13, 1011–1021.

(41) Souaille, M.; Roux, B. Extension to the weighted histogram analysis method: combining umbrella sampling with free energy calculations. Computer Physics Communications 2001, 135, 40–57.

(42) Essmann, U.; Perera, L.; Berkowitz, M. L.; Darden, T.; Lee, H.; Pedersen, L. G. A smooth particle mesh Ewald method. The Journal of Chemical Physics 1995, 103, 8577–8593.

